# A transition state-like acylenzyme conformation distinguishes carbapenemase activity in class A β-lactamases

**DOI:** 10.64898/2026.08.31.748333

**Authors:** Michael Beer, James Spencer, Adrian J. Mulholland

## Abstract

Carbapenems are the most potent β-lactams, key antibiotics for healthcare-associated infections by Gram-negative bacteria and evade hydrolysis by most β-lactamases; but are increasingly threatened by emergence of enzymes exhibiting hydrolytic activity towards them. Of the four recognised β-lactamase subclasses, class A (active-site serine enzymes that hydrolyse β-lactams via a covalent acylenzyme intermediate) is the most widely disseminated and, while the majority of such enzymes react with carbapenems to form long-lasting acylenzyme complexes, several possess carbapenem-hydrolyzing activity (carbapenemases). Here, we investigate the basis for these differences in a panel of class A beta-lactamases using molecular dynamics (MD) simulations of the respective acylenzyme complexes and tetrahedral intermediates (TI). The simulations reveal multiple features associated with catalytic activity across the spectrum of enzymes tested, including more extensive interactions of the carbapenem acylenzyme carbonyl and generally increased lifetimes of active site water molecules positioned for deacylation. Analysis of the dynamic trajectories shows carbapenemases to have reduced root mean-squared fluctuation (RMSF) differences between the acylenzyme and TI, that are not limited to the active site, indicating that the acylenzyme complex is pre-organised for reaction in carbapenemases but not in carbapenem-inhibited enzymes. Similarly, Principal Component Analysis (PCA) of acylenzyme and TI dynamics shows greater overlap between the two states in carbapenemases, providing further evidence for acylenzyme pre-organisation. Such simulations may represent an effective computational assay able to identify enzymes with carbapenemase activity at relatively modest computational cost.

## Introduction

The spread of antimicrobial resistance (AMR) has become one of the greatest threats to medicine and human health (1). Currently available antibiotics are becoming increasingly ineffective in treating bacterial diseases, with England reporting numbers of bacteraemia episodes, caused by one of *Acinetobacter* spp., *Escherichia coli*, *Enterococcus* spp., *Klebsiella oxytoca, K. pneumoniae, Pseudomonas* spp., *Staphylococcus aureus* or *Streptococcus pneumoniae* that are resistant to one or more antibiotics, increasing by 3.5 % between 2019 and 2023 (2). On the current trajectory, by 2050 AMR will result in 10 million excess global deaths per year, along with a loss of $100 trillion in productivity (1, 3). In response to this growing threat the Infectious Diseases Society of America expressed the pressing need for AMR research and the development of novel antimicrobial therapies (4).

β-Lactams constitute 65% of all antibiotic prescriptions (5) and are the most important class of antibacterial therapeutics. The four major β-lactam classes: penicillins, cephalosporins, carbapenems and monobactams; differ in the functional groups attached to the β-lactam ring (5). Later-generation cephalosporins and carbapenems are the most recent β-lactam antibiotics to enter the clinical market, with carbapenems initially described as ‘last resort’ antibiotics, used to treat Gram-negative bacterial infections showing resistance to many/all other antibiotics (6). β-lactams are hydrolysed by bacterial enzymes known as β-lactamases. These enzymes represent the primary β-lactam resistance mechanism in Gram-negative bacteria (7). The evolution and dissemination (on plasmids and other mobile genetic elements) of β-lactamases has accelerated in response to the widespread use of β-lactam antibiotics (8).

The Ambler classification system divides β-lactamases into four classes (A, B, C and D), based on structural and sequence similarity (9). Classes A, C and D are the serine β-lactamases (SBLs), that utilise a hydrolysis mechanism involving nucleophilic attack by an active site serine, forming an acylenzyme intermediate (10). This is subsequently deacylated by addition of a water molecule to generate a hydrolysed product, devoid of antibiotic activity. Class B β-lactamases contain 1 or 2 active site zinc ions that activate a water molecule for nucleophilic attack on the scissile β-lactam amide. Class A is the most widely disseminated of the four Ambler classes and is distinguished by the conserved, catalytically important active site residue Glu166 (10). Carbapenem hydrolysis by class A β-lactamases is a two-step process: the first step, acylation, involves nucleophilic attack of the activated Ser70 hydroxyl on the carbonyl amide carbon (C7) of the β-lactam ring, forming the acylenzyme complex with concomitant cleavage of the β-lactam amide. The subsequent deacylation step then involves proton transfer from the deacylating water molecule (DW) to Glu166, followed by nucleophilic attack on C7, resulting in hydrolysis of the acyl bond via a tetrahedral intermediate (Scheme 1). Slow deacylation rates are often the cause of carbapenem inhibition of class A β-lactamases (10).

**Scheme 1:**
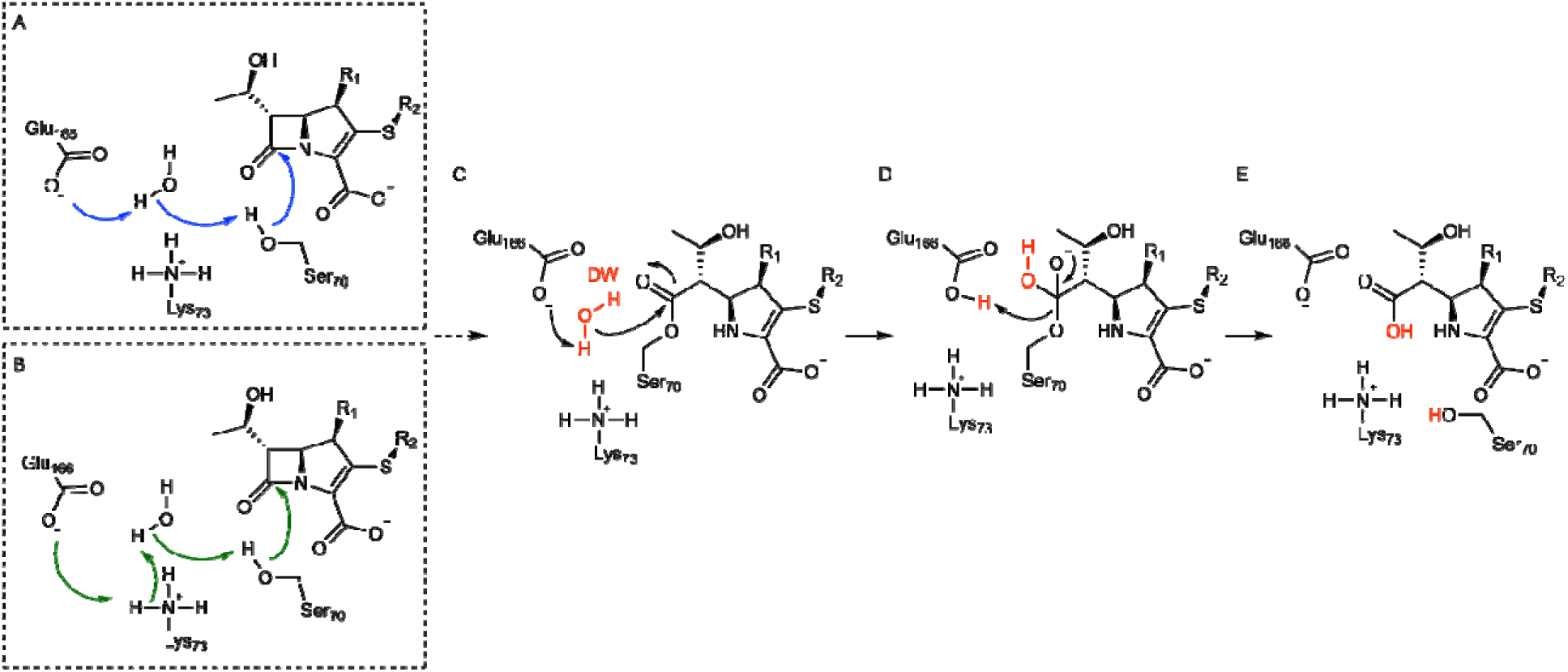
A general mechanism of carbapenem hydrolysis by class A β-lactamases. The acylation pathway can take of two mechanisms (A or B) resulting in an acylenzyme complex (C). This then undergoes deacylation, going through the deacylation tetrahedral intermediate (TI, D), where a deacylating water (DW, red) has a proton abstracted by Glu166, activating it for nucleophilic attack. This results in a hydrolysed product (E), devoid of antibiotic activity.

Catalytic activity against a broad range of clinically available β-lactams is being detected with increasing regularity, especially amongst Class A enzymes (11–15). Of concern are the increasing numbers of extended-spectrum β-lactamases (ESBLs, that hydrolyse oxyiminocephalosporins) and carbapenem-hydrolysing β-lactamases (carbapenemases) being detected in the clinic (15–17). The U.S. Centers for Disease Control (CDC) has recognised the risk that both ESBLs and carbapenemases pose by categorising ESBL-producing and carbapenem-resistant Enterobacterales as Serious and Urgent Antibiotic Resistance Threat pathogens, respectively (18).

Understanding the mechanisms by which class A β-lactamases can hydrolyse carbapenem antibiotics is vital for development of the carbapenem scaffold to generate novel therapeutic agents that evade this resistance mechanism. Previous computational studies by our group have used quantum mechanics/molecular mechanics (QM/MM) simulations to study the carbapenem deacylation reactions of class A β-lactamases, identifying structural features that promote carbapenemase activity (19, 20). QM/MM simulations, although superior in their ability to provide accurate descriptions of the geometries of molecules within their QM regions, suffer from limited sampling with currently available computational resources. Hence, long timescale molecular mechanics (MM) molecular dynamics (MD) simulations may provide further insight into determinants of carbapenemase activity, particularly informing on the dynamic ensembles of the acylenzymes of carbapenemases and comparator enzymes that lack carbapenemase activity, and against which carbapenems function as effective covalent inhibitors.

The deacylation reaction progresses through a tetrahedral intermediate (TI, Scheme 1), the stability of which is known to be important (19). Our previous work found that electric field analysis, particularly of the TI ensemble, can effectively and significantly discriminate between class A β-lactamases that can and cannot hydrolyse carbapenems(20). Other studies too have highlighted the importance of stabilising the TI in a range of enzyme systems (21–23). Understanding the dynamic ensemble of this species may help describe transition state-like complexes of class A β-lactamases, ultimately enabling development of transition state-like inhibitors, a strategy that has already proved fruitful as an approach to countering β-lactamase activity (24). Furthermore, these works highlight dynamical changes between a ground state ensemble, such as the acylenzyme complexes presented here, and that of a high energy intermediate, such as the TI. This may provide greater understanding of factors defining efficient catalysis, such as in the β-lactamase mediated hydrolysis of carbapenems.

Accordingly, we here present a total of 1.5 μs of MM MD simulation of the acylenzymes formed on reaction of the carbapenem meropenem with each of the 8 β-lactamases previously studied with QM/MM simulations (20, 25); along with a further 1.5 μs of MM MD simulations of each of the meropenem-derived deacylation TI, totalling 24 μs of simulation time. We analyse the dynamic ensembles of both species, examining the differences between the acylenzyme and the TI. We show that the carbapenemase enzymes exhibit similar dynamics in both states, suggesting that a transition state-like conformation is frequently sampled in the acylenzyme, whereas acylenzymes of carbapenem-inhibited enzymes sample a range of conformations that differ from the TI ensemble, and are thus likely catalytically incompetent. We further analyse, across the panel of enzymes, the electrostatic stabilisation of the acylenzyme and the TI and the hydration of the active site. Overall, the simulations presented here highlight how the dynamic ensembles sampled by carbapenem-inhibited enzymes in the acylenzyme and the TI differ from those sampled in carbapenemases; thus, indicating how acylenzyme dynamics is a determinant of carbapenemase activity.

## Methods

### Simulation System

For the simulations of the meropenem-derived acylenzymes, the starting structures and simulation systems were those used in previous publications (19, 26–28), except for KPC-2 (see below). The BlaC, SHV-1 and SFC-1 starting structures were obtained from crystal structures of the respective meropenem complexes (PDB IDs 3DWZ (29), 2ZD8 (30) and 4EV4 (31)). For TEM-1, meropenem was modelled into the electron density observed for bound imipenem in the crystal structure of the TEM-1-complex (PDB ID 1BT5 (32)). CTX-M-16 (PDB ID 1YLW (33)), NMC-A (PDB ID 1BUE (34)), and SME-1 (PDB ID (35)) uncomplexed structures were aligned with the crystal structure of the SFC-1:meropenem complex, and meropenem modelled into the respective active sites. For KPC-2 the meropenem-derived acylenzyme structure (PDB ID 8AKL (36)) was used with the meropenem Δ2 tautomer modelled into the active site. In all cases structural alignments and meropenem modelling was undertaken using Wincoot (37) and the secondary structure matching (SSM) superpose algorithm (38). As previously described, GAFF was used to parameterise acylated meropenem, with partial charges derived using the R.E.D. server (27, 39).

For simulations of the TI complexes, snapshots that correlated with the TI in previous QM/MM investigations, as described by Jabeen *et al* (28), with the solvent atoms removed, were taken as starting structures. For KPC-2, the meropenem-derived acylenzyme structure ((PDB ID 8AKL (36)) was aligned to a snapshot of the TI complex taken from simulations in Jabeen *et al.,* (20) (in short, this involved aligning uncomplexed KPC-2 (PDB ID 2OV5) to the SFC-1:meropenem acylenzyme structure (PDB ID 4EV4) and using the adaptive string method (40) to model formation of the TI) and the meropenem TI structure merged into the new KPC-2 enzyme structure. Meropenem TI parameters were constructed using the values calculated from a previously published serine hydrolase TI parameter set and GAFF parameters, after geometry optimisation and partial charge derivation using the R.E.D. server (39, 41, 42).

Systems were hydrated in a solvent box of TIP3P water, to a distance of 10 Å away from any solute atoms. Na+ and Cl-ions were added to neutralise the charge of the system. Protonation states of all ionisable residues were determined using PROPKA at pH 7.4 (43).

### Minimisation, Heating and Equilibration

All minimisation, heating and equilibration steps were performed using the AMBER20 package (44). Each system was minimised for 1000 steps, with 300 steps of steepest descent and 700 steps of conjugate gradient and 500 kcal/mol.Å^-2^ harmonic restraints on all backbone atoms. The minimised system was then heated from 25 K to 300 K in the NVT ensemble over 50 ps, with 100 kcal/mol.Å^-2^ restraints on backbone atoms, a long-distance electrostatic cutoff of 10 Å, and using the Berendsen thermostat with a coupling constant of 1.0 ps and a collision frequency of 1 ps^-1^. Four steps of equilibration in the NPT ensemble were then performed at 300 K and a pressure of 1 atm (coupling constant 2 ps). Each equilibration step was 250 ps with a 10 Å long-range electrostatic cutoff. Restraints on Cα atoms were reduced from 50 kcal/mol.Å^-2^ to 0 kcal/mol.Å^-2^ over the course of the 1 ns (over four steps of 250ps) of equilibration.

### Production MD Simulations

Production simulations, run using the AMBER20 package (44) for both the acylenzymes and the tetrahedral intermediates, were performed in the NPT ensemble, with a 2.0 ps coupling constant for pressure and temperature. Long-range electrostatic cutoff was set at 10.0 Å and simulations were run for 500 ns. Three repeats each were performed for the acylenzymes and tetrahedral intermediate of each enzyme, resulting in 3 μs of production simulation per enzyme. Root mean square deviation (RMSD) analysis was used to confirm the production simulations were stable (Figures S1, S2).

### Analysis

Root mean square fluctuation (RMSF), distance and hydrogen bonding analyses were performed using the CPPTRAJ module of AMBER20 (44). The MDAnalysis python module was used to analyse survival probability of the deacylating water molecules and principal component analysis (45).

## Results

In previous studies (20, 25, 46) we have used QM/MM approaches to investigate deacylation of the meropenem acylenzymes of 8 representative class A β-lactamases, that differ in their carbapenem-hydrolysing activity. KPC-2, NMC-A, SME-1 and SFC-1 are all carbapenemases, whereas TEM-1, SHV-1, CTX-M-16 and BlaC are all inhibited by carbapenems. We here extend these investigations to examine the dynamic behaviours of two mechanistically relevant species, the meropenem acylenzyme and tetrahedral oxyanion intermediate (TI) formed during the deacylation reaction, over longer (μs) time scales using MM MD simulations. Specifically, we interrogate the simulations to investigate differences between the respective acylenzyme and TI ensembles. In particular, we investigated the electrostatic stabilisation of the meropenem-derived carbonyl; the orientation in the active site of the meropenem-derived species; the positional stability of water molecules positioned for deacylation; and the dynamic behaviour of each enzyme, focusing on the mobile loops adjoining the active site and how their behaviours change between the acylenzyme and tetrahedral intermediate states.

Analysis of simulation trajectories (three 500 ns simulations for the meropenem acylenzyme and deacylation tetrahedral intermediate (TI) for each enzyme studied, Figures S1, S2) demonstrates convergence of all the simulations, as evidenced by the time-dependence of Cα RMSD values. Of note, for KPC-2 the N- and C-termini are particularly flexible, resulting in higher RMSD values; an observation we have made in previous studies (47). Each simulation was visually checked to confirm that the higher RMSD values were due to fluctuations in the N- and C-termini rather than large conformational changes in the core residues. On this basis, we then focused on specific properties of the respective systems, seeking to discriminate between enzymes differing in activity towards carbapenems.

### The Acylenzyme and Tetrahedral Intermediate Make Persistent Hydrogen Bonds to the Oxyanion Holes of Carbapenemase Enzymes

During the deacylation reaction, a negatively charged oxyanion is formed (Scheme 1, Figure S3) (48) that is stabilised by the oxyanion hole formed by the backbone amide nitrogen atoms of residues at positions 70 and 237. Previous studies have determined that, in some class A β-lactamases that lack carbapenem-hydrolysing activity, the acylenzyme carbonyl oxygen (Figure S4) can sample an orientation ‘outside’ of this oxyanion hole, leading to suggestions that effective stabilisation of carbapenem-derived oxyanions may be a point of difference between class A carbapenemases and related enzymes lacking carbapenem-hydrolysing activity (30, 32). In the simulations presented here, all of the starting structures positioned the carbonyl oxygen atoms of the respective meropenem-derived species within the oxyanion hole, as was the case for our previous QM/MM simulations (19, 28); rather than in the “flipped-out” conformations observed in crystal structures of TEM-1 and SHV-1 carbapenem derived acylenzymes (30, 49). However, distance analysis between the oxyanion and the Ser70 backbone amide showed that in the meropenem-derived CTX-M-16 acylenzyme the oxyanion transiently “flipped” outside of the oxyanion hole during one simulation, increasing the distance between the two atoms to more than 5 Å (Figure S5). This was not, however, observed in the trajectories of any other meropenem derived acylenzymes, including TEM-1 (PDB ID 1BT5 (32)) and SHV-1 (PDB ID 2ZD8 (30)) where “flipped-out” conformations have been observed in some crystal structures of carbapenem derived acylenzymes (30, 49).

More recently, we have shown, using QM/MM molecular dynamics simulations, that the nature of the electric field acting upon the oxyanion carbonyl discriminates between the two groups of enzymes (28). Based on these findings, we analysed hydrogen-bonding interactions of the two components of the oxyanion hole (the Ser70 and Ala/Thr/Ser237 backbone amides) over the 1.5 μs of MM MD simulation for the meropenem acylenzyme and tetrahedral deacylation intermediate of each enzyme (Figure 1). This aimed to test whether direct electrostatic interactions, in both the acylenzyme and tetrahedral intermediate ensembles, differ between carbapenemases and enzymes lacking carbapenemase activity in our set of representative β-lactamases.

**Figure 1.**
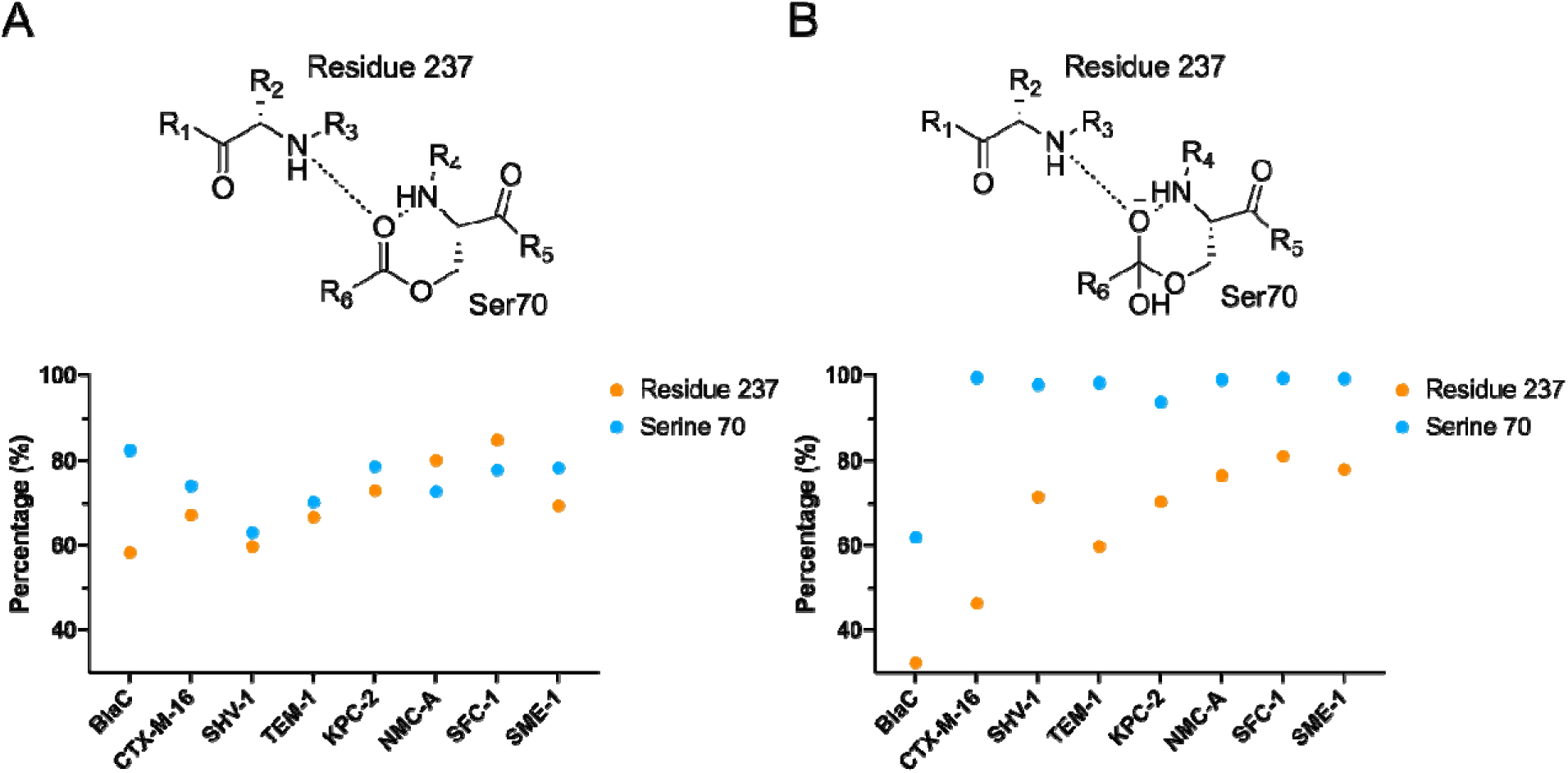
Hydrogen Bonding Interactions in the Oxyanion Holes of the Acylenzyme and Tetrahedral Deacylation Intermediate. Plots show percentage of frames in which a hydrogen bond is formed between the backbone amides of Ser70 and residue 237 to the carbonyl oxygen of meropenem-derived species in A) the acylenzyme and B) the tetrahedral deacylation intermediate.

This analysis indicates that, in the meropenem-derived acylenzymes, the backbone amide of residue 237 more consistently makes hydrogen bonds to the carbonyl oxygen (Figures 1, S4) in carbapenemases than in carbapenem-inhibited enzymes. In SHV-1 and TEM-1, both carbapenem-inhibited enzymes, the backbone amide of serine 70 also makes less consistent hydrogen bonds to the acylenzyme carbonyl group than in any carbapenem-hydrolysing enzyme. However, in the BlaC and CTX-M-16 enzymes, which also lack activity towards carbapenems, the extent of hydrogen bonding involving the serine 70 backbone amide approaches that observed in the carbapenemases. In the simulations of the tetrahedral deacylation intermediates (TI), the carbapenem-inhibited enzymes BlaC, CTX-M-16 and TEM-1 all form less stable hydrogen bonds between residue 237 and the oxyanion than do the carbapenem-hydrolysing enzymes. In addition, in simulations of the TI BlaC also makes relatively few hydrogen bonds between the Ser70 backbone amide and the oxyanion. These analyses suggest that, with respect to hydrogen-bonding interactions to the carbonyl oxygen of meropenem-derived species, in the acylenzyme and in the TI, the contributions of the backbone amide of residue 237 can partially discriminate between the carbapenemase and carbapenem-inhibited enzyme groups.

The presence of an active site disulfide bridge, between residues Cys69 and Cys238 (neighbouring Ser70 and residue 237 that form the oxyanion hole) is a noted feature of Class A carbapenemases (50). To test whether the reduced hydrogen bonding to the acylenzyme carbonyl oxygen observed in carbapenem-inhibited enzymes is due to the lack of this disulfide bridge, we analysed the distance between the Cα atoms of residues 69 and 238 in all tested enzymes. An increase in this distance would indicate that that the active site is less compact in the vicinity of the backbone amides of Ser70 and residue 237 (i.e the oxyanion hole), possibly explaining the observed reduction in hydrogen bonding. The results show that in the meropenem acylenzymes formed by the carbapenem-inhibited enzymes, which lack this disulfide bond, the average distance between residue 69 and residue 238 was greater (Table S3).

### Acylated Meropenem adopts a distinct orientation in the active site of Class A Carbapenemases and the *M. tuberculosis* β-lactamase BlaC

The orientation of the bound carbapenem ligand within the active site is likely to have a significant effect upon its susceptibility to hydrolysis. To test whether bound meropenem adducts adopt different conformations in the active sites of class A carbapenemases, compared with enzymes that do not hydrolyse carbapenems, we analysed the hydrogen bonding interactions between each of the eight proteins and their respective meropenem-derived adducts.

Due to sequence differences between class A carbapenemases and those lacking carbapenemase activity, the meropenem derived acyl-adducts make different hydrogen bonding interactions with residues within the enzyme active site. Of note, the carbapenem-inhibited enzymes CTX-M-16, SHV-1 and TEM-1 all form hydrogen bonding interactions between the meropenem C3 carboxylate and the side chains of residues Arg244 or Arg276. The four carbapenemase enzymes, as well as BlaC, feature either Aspartate or Glutamate at these positions (Figure S6). Furthermore, in SHV-1 and TEM-1, the meropenem C3 carboxylate makes hydrogen bonds with the invariant active site residues Lys234 and Ser130 (Figure 2). This results in the meropenem-derived covalent adduct adopting a different orientation within the active sites of CTX-M-16, TEM-1 and SHV-1, compared to the carbapenem-hydrolysing enzymes, or BlaC (Figure 2).

**Figure 2.**
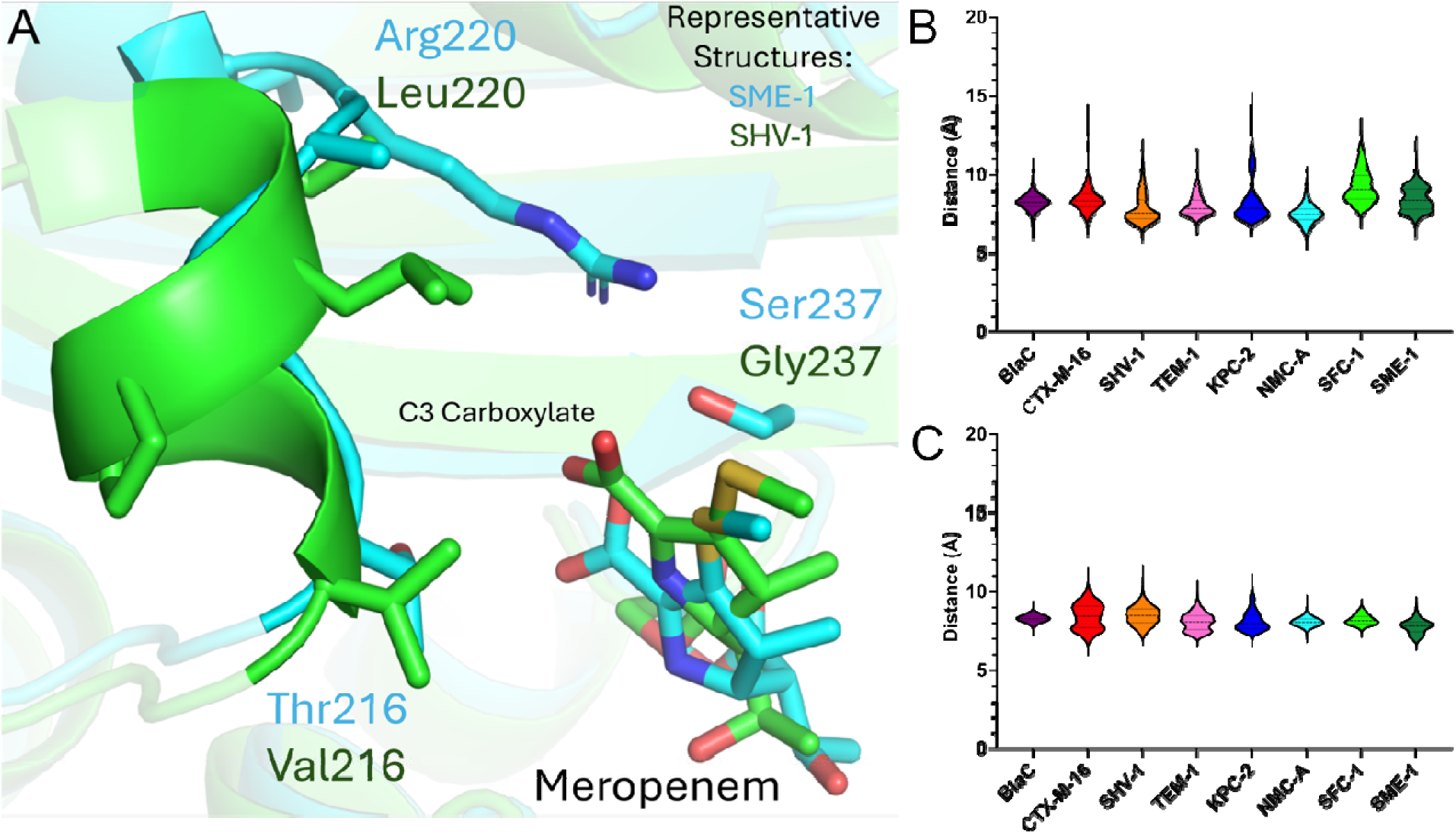
Distance of bound carbapenem from beta-lactamase hinge region. A) Representative conformation from the SME-1 and SHV-1 trajectories highlighting the differences in meropenem acylenzyme orientation between SME-1 (carbapenemase) and TEM-1 (carbapenem-inhibited) enzymes. B) Violin plot showing the distribution of distances of the centre of mass of residues 216-220 to the C3 carboxylate of meropenem in the acyl-enzyme and C) TI.

The carbapenemase enzymes, along with BlaC, all contain an arginine residue at Ambler position 220 (part of the “hinge region” formed by residues 213 – 220), where the other non-carbapenemase enzymes have a leucine (Figure S6). In BlaC and SME-1 this arginine remains consistently hydrogen bonded to the C3 carboxylate of bound meropenem-derived species throughout the simulations of both the acylenzyme and TI (Figure 2). In NMC-A, Arg220 remains hydrogen bonded to the meropenem C3 carboxylate for smaller proportions of the simulations of the acylenzyme (< 15 %) and TI (< 45 %). Alternatively, in simulations of KPC-2 and SFC-1, Arg220 is hydrogen bonded to the side chain of Ambler position 237 (which is proximal to the C3 carboxylate of bound adducts, Table S1) for 100% of the frames for both enzymes in the acylenzyme and TI. The consequence of either of these interactions is that the loop forming the hinge region (residues 213-220) remains close to the C3 carboxylate (Figures S7, S8) apparently helping to maintain the orientation of meropenem-derived species. These interactions are less pronounced in the simulations of the carbapenem-inhibited enzymes, due to the lack of hydrogen bonding involving residues 220 and 237, resulting in simulations regularly sampling a different orientation of the meropenem-derived covalent adduct, compared to carbapenemase enzymes (Figure 2).

One effect of these differences in orientation may be upon the distance between the β-lactam amide nitrogen (N4) and the carbonyl oxygen, potentially impacting electrostatic stabilisation of the TI oxyanion (19, 20, 46). To explore this possibility, we analysed the distance between the carbonyl oxygen and the β-lactam amide in the simulations of the various TI and acylenzyme complexes. In simulations of the meropenem-derived acylenzymes, the distances between these two atoms were significantly greater (confirmed by pairwise student t-tests, Table S3) for TEM-1 and SHV-1 than for any of the carbapenem-hydrolysing enzymes (Figure S8). However, in the simulations of the TI, only for BlaC was this distance significantly greater than in the carbapenemase enzymes.

### Carbapenemases Effectively Stabilise a Water Molecule in the Deacylating Water Position

Appropriate positioning of a water molecule in the deacylating position (DW) is essential to efficient deacylation of carbapenem acylenzymes, due to the requirement for the DW to be activated by Glu166 to enable nucleophilic attack upon the C7 carbon (36, 51, 52). A survival probability script, written using the MD Analysis package (45), was written to determine the likelihood of a water molecule remaining in the deacylating position over time during simulations of the respective carbapenem acylenzymes.

Analysis of the survival probability of bound DW for all 8 β-lactamases tested shows that, in the meropenem acylenzymes of BlaC and CTX-M-16, there is a reduced probability of a water molecule remaining in a position compatible with activation by Glu166 for more than 200 ps, compared to the other enzymes. At 1 ns, the chance of a water molecule being retained in the deacylating position is 16.7 % and 5.8 % for BlaC and CTX-M-16, respectively, whereas for the remaining enzymes this probability is between 35.6 - 53.7 %. Notably, however in simulations of the acylenzymes of the carbapenem-inhibited enzymes SHV-1 and TEM-1, the survival probabilities of water molecules in the deacylating position (49.5 % and 47.4 %, respectively, after 1 ns) are similar to those observed for the carbapenemases.

### Carbapenem Acyl-Enzymes of Carbapenemases Frequently Sample Transition State-Like Conformations

Acylenzymes of class A β-lactamases can sample a range of conformations, many of which are catalytically inert (53). Pre-organisation of a ground state is known to be an important factor for efficient enzyme-mediated catalysis. A transition-state-like conformation, as defined by Walker *et al.* (54), describes conformations of the ground-state (in this case the acylenzyme) ensemble equivalent to those found in the transition state ensemble (here the analogous TI). To investigate the presence of transition-state-like conformations in meropenem acylenzymes of carbapenem-inhibited and carbapenem-hydrolysing enzymes we analysed the dynamical behaviour of both the acylenzyme and TI ensembles, for all 8 enzymes, using root mean square fluctuations (RMSF).

The differences in the RMSF (ΔRMSF) between the TI and the acylenzyme were measured to determine changes in fluctuation. The carbapenem-inhibited enzymes exhibited consistently low ΔRMSF values across the entirety of the enzyme structure. In contrast, the carbapenem-inhibited enzymes BlaC, SHV-1 and TEM-1 exhibited large differences in fluctuations between the acyl-enzyme and the TI for the catalytically important Ω-loop (residues 164-179, Figure S9). In particular, this apparently reflects increased flexibility of the Ω-loop in the BlaC and SHV-1 acylenzymes (Figures S9-11). In addition, the 2β4 loop (residues 87 – 94), previously shown to be important in the catalysis of class A β-lactamases, exhibits large decreases in fluctuation on progression of the acylenzyme to the TI state for the carbapenem-inhibited BlaC, CTX-M-16 and TEM-1 enzymes. The carbapenemases and SHV-1 show little change or an increase in flexibility in this loop in the TI compared to the acylenzyme (Figure S9). The 3 4 loop (residues 102 – 108), which includes residue 105, known to be important in defining catalytic conformation (55), also shows large changes in flexibility between the acylenzyme and the TI in the carbapenem-inhibited BlaC and TEM-1 enzymes, with notably large changes in the dynamic behaviour of TEM-1 residue 105. Of note, residue 104, previously identified as a residue that can modulate catalysis, fluctuates much more in the CTX-M-16 TI than the respective acylenzyme.

The 240-loop (residues 237 - 244) previously identified as being catalytically important in KPC-2, undergoes a flexible to rigid transformation when progressing from the acylenzyme to the TI for all four carbapenemases investigated. In particular, KPC-2 Val240 undergoes extensive rigidification in the TI ensemble relative to the acylenzyme. CTX-M-16, displays the opposite behaviour, with multiple residues in the 240-loop increasing in flexibility when compared with the acylenzyme (Figure 3, S9).

**Figure 3.**
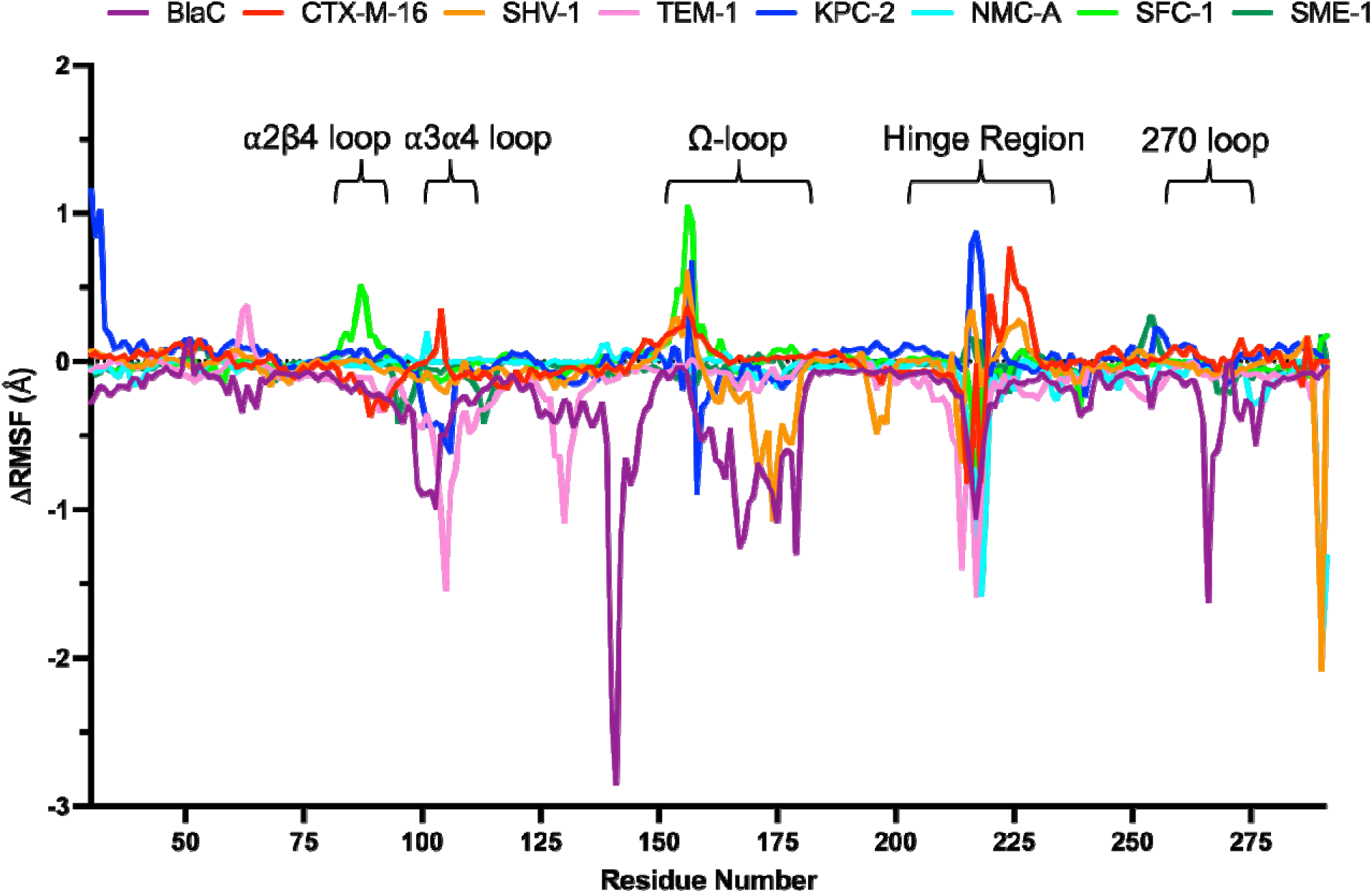
Changes in residue fluctuations between the acylenzyme and TI ensembles. Differences in root mean square fluctuation values between simulations of the TI and acylenzyme. Negative values indicate that the acylenzyme has a greater RMSF value (i.e. undergoes more extensive fluctuations of Cα atoms) than the TI; whilst positive values indicate that the TI has a greater RMSF value than the acylenzyme. In all cases Cα fluctuations are calculated based on comparisons with the minimised initial structures.

To further examine the global changes in dynamics, we undertook principal component analysis (PCA) of the C RMSF data in the simulations of both the acylenzyme complexes and the TI for all eight enzymes (Figure S12). Visual inspection of each plot showed high overlap between the acylenzyme complexes and TI for the carbapenemase enzymes, whilst this was less pronounced for the carbapenem-inhibited enzymes. This is supported by overlap and Bhattacharyya coefficient analysis (Figure 4, Table S4), interrogating the similarity of the acylenzyme complex and TI PCA projections, whereby all carbapenem inhibited enzymes had an overlap coefficient and Bhattacharyya coefficient score of less than 0.25 and 0.5, respectively, whilst all carbapenemases had scores above these values (Table S4). Clustering of the PCA plots indicated that, for three of the four carbapenemase enzymes, the most highly populated cluster for the acylenzyme complex matched the most populated cluster in the TI projection. The exception was SME-1, which showed a more even distribution across all three clusters in both the acylenzyme complex and TI states (Table S5). Carbapenem-inhibited enzymes, other than BlaC, showed the opposite, with the most populous clusters in the acylenzyme complex being (among) the least populous clusters in the TI. Whilst BlaC had similarly populous clusters between the acylenzyme and TI, the overlap between projections is low, indicating that there were still differences between the two ensembles and that BlaC does not regularly sample catalytically competent conformations.

**Figure 4.**
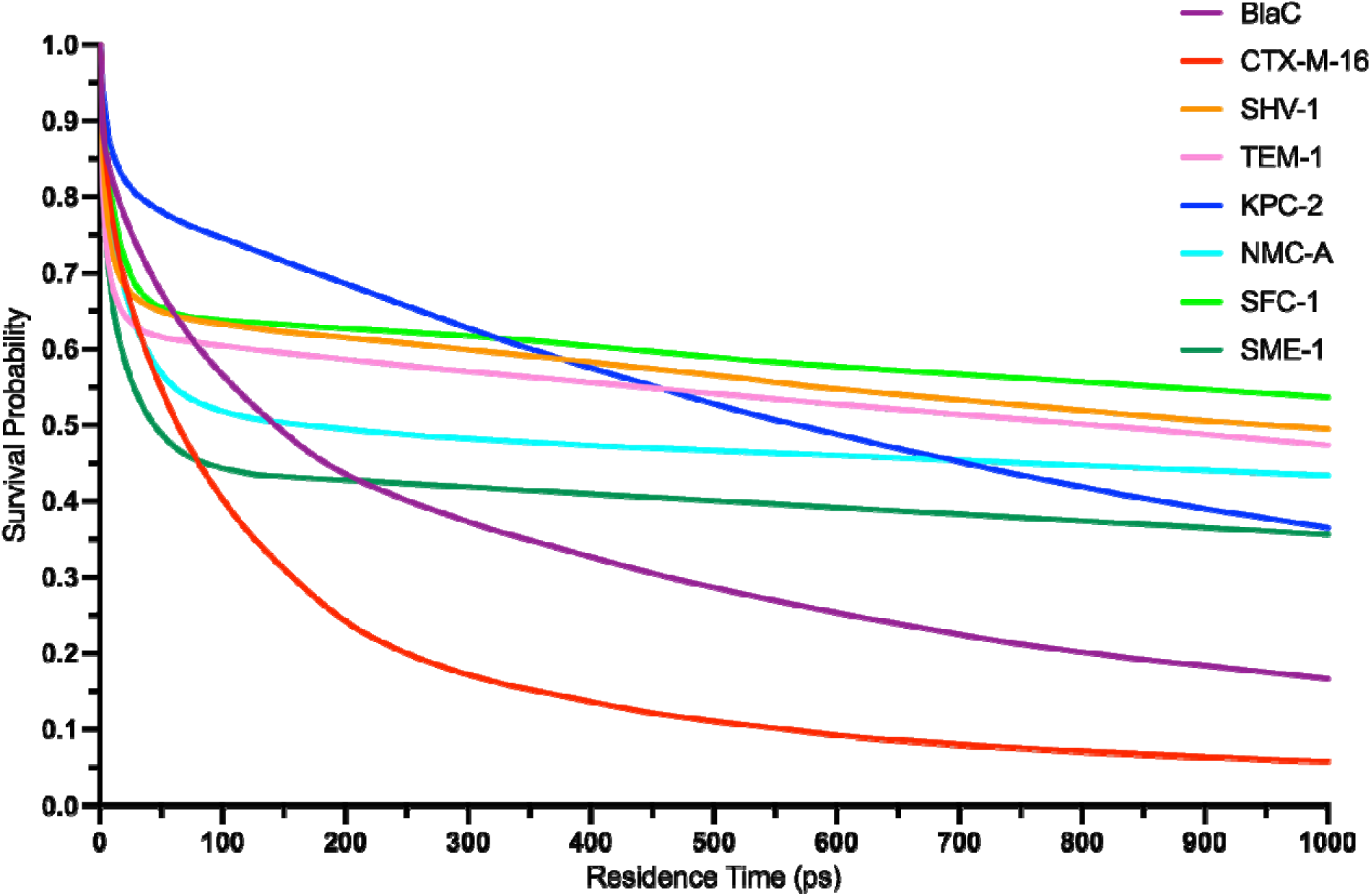
Survival Probability of a Water Molecule Remaining in the Deacylating Water Position over Time. The probability of a water molecule remaining in the deacylating position (defined as a sphere with a radius of 3.5 Å in the geometric centre of the Glu166^Cδ^ and the carbonyl carbon of the meropenem derived acyl adduct) over time, upon entering the deacylating water position.

**Figure 4.**
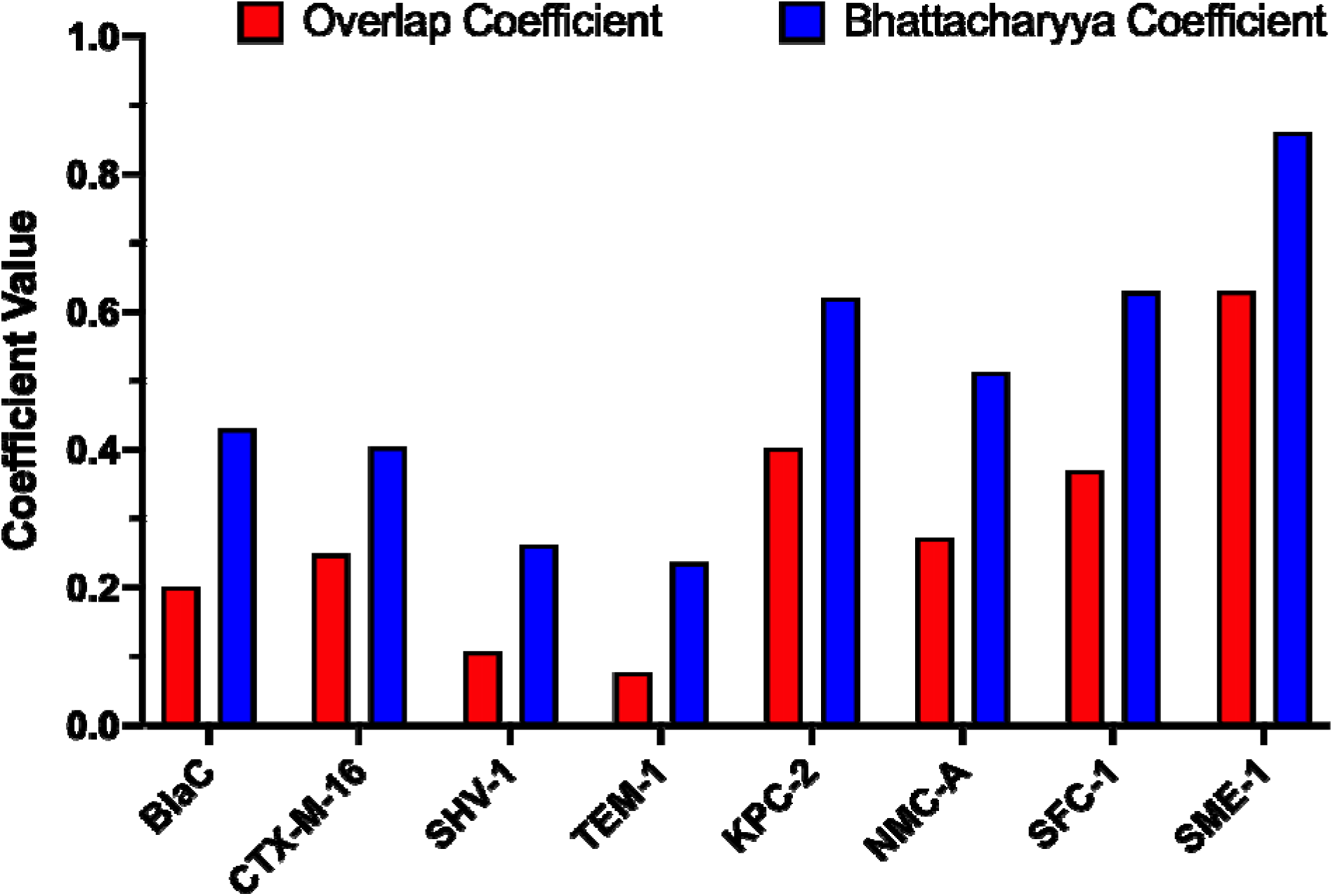
Comparison of the PCA projections across the eight-class. **A** β**-lactamases.** For each enzyme the overlap coefficient (red) and the Bhattacharyya coefficient (blue) are shown.

## Discussion

This work has taken a representative group of four class A β-lactamases that can efficiently hydrolyse carbapenems, and a further four class A enzymes that cannot, and subjected these to a total of 24 μs of molecular dynamics simulation, involving simulations of the respective acylenzyme complexes and deacylation tetrahedral intermediates (TI) of the carbapenem meropenem. Previously published QM/MM studies of these enzymes have demonstrated that the carbapenemase and carbapenem-inhibited groups could be distinguished by comparing the calculated free energy barriers for TI formation, and analysing the active site electric field in the deacylation transition state or TI (19, 28). Here we analyse the dynamic ensembles of both the acylenzymes and tetrahedral intermediates, that collectively provide greater understanding of why a select few class A β-lactamases can effectively catalyse hydrolysis of carbapenem antibiotics, whereas others cannot.

Electrostatic stabilisation of substrates bound in the active site is a recognized feature of enzyme-catalysed reactions and is generally agreed to be a requirement for efficient catalysis (28, 56, 57). Deacylation of β-lactamase acylenzymes of carbapenem antibiotics progresses through an oxyanionic TI (10). Effective stabilisation of the oxyanion during the deacylation reaction has been proposed as one feature that may discriminate between carbapenemases and enzymes that lack carbapenemase activity (28, 30, 32, 58). Hydrogen bonding analysis, presented here, of the interactions involving the oxyanion hole in the studied enzymes indicates that carbapenemases support stronger electrostatic interactions with bound carbapenem-derived species, through hydrogen bonds to the oxyanion hole, in both the acylenzyme and TI. This increased electrostatic stabilisation of the carbonyl oxygen is largely determined by differences in hydrogen bonding involving the backbone amide of residue 237, corroborating studies showing a reduction in carbapenemase activity in KPC-2 and SME-1 when this residue is mutated (59, 60), and appears to be a direct result of the additional active site disulfide bond (between Cys69 - Cys238) that is a feature of class A carbapenemases. In all carbapenemases tested, the distance between residues 69 and 238 is consistently smaller than is the case in the comparator enzymes, due to the restraints imposed by the disulfide bond. Indeed, this restricts the distance between the proximal residues 237 and Ser70 in the carbapenemases, compared to the carbapenem-inhibited enzymes, particularly in the TI. This feature would likely stabilise both the transition state between the acylenzyme and the TI, and the TI itself, so promoting the deacylation reaction. Our previous QM/MM simulations of an SFC-1^C238G^ mutant reveal that disruption of this disulfide bridge results in an increased barrier to deacylation (19). It is further noted that there is a general increase in the extent of hydrogen bonding of the carbonyl oxygen to the oxyanion hole in the TI, compared with the acylenzyme, particularly in carbapenemase enzymes. This implies that for efficient carbapenemase activity the portions of the protein that form the oxyanion hole sample a restricted dynamical landscape in the TI compared with the acylenzyme, resulting in increased hydrogen bonding to the carbonyl oxygen. This is consistent with later RMSF analysis, which identified a general reduction in residue fluctuations in carbapenemases, compared with non-carbapenemases.

We note that some of the GES-type enzymes, not investigated here, possess the equivalent disulfide but lack activity towards carbapenems (61). This suggests that presence of the Cys69 – Cys237 disulfide is not the sole factor determining whether the carbapenem-derived oxyanion can be sufficiently stabilised to achieve carbapenemase activity. Consistent with this reasoning, a C69G mutation of the GES-5 enzyme, which can hydrolyse carbapenems, reduces, but does not eradicate, carbapenemase activity.

Analysis of the crystal structures of GES-5 and GES-5^C69G^ has indicated that the C69G mutant contains a hydrogen bonding network that ensures the compactness of the oxyanion hole, so explaining the maintenance of carbapenemase activity in the absence of a disulfide bond between residues 69 and 238 (50).

Differing hydrogen bonding networks in the groups of carbapenem-hydrolysing and carbapenem-inhibited enzymes, together with differing conformations of the hinge region loop (residues 213 – 220), can also influence activity towards carbapenems via effects upon the orientation of the carbapenem derived acyl-adduct, as observed in CTX-M-16, SHV-1 and TEM-1. This can be characterised most simply as differences in the orientation of the C3 carboxylate, in the carbapenemase enzymes resulting in enhanced electrostatic interactions between the β-lactam carbonyl oxygen and amide nitrogen in the respective acylenzyme adducts, evidenced as smaller distances between these two atoms (Figure 2). This provides additional stabilisation of the charge that accumulates during the transition to the oxyanionic TI following nucleophilic attack of the deacylating water molecule upon the acylenzyme carbonyl. Such differences in orientation of the C3 carboxylate have been observed previously, for example in meropenem acylenzyme adducts of the KPC-2 carbapenemase. In that case the C3 carboxylate of the Δ1-2*R* acylenzyme tautomer, that is considered to be deacylation-incompetent, adopts a similar orientation to that described here for meropenem-derived species bound to carbapenem-inhibited enzymes. (Note that in this work all meropenem-derived species are modelled in the Δ2 tautomer and alternative configurations are not considered). The orientation observed for KPC-2 was thought to provide less effective electrostatic stabilisation, compared to the deacylation-competent Δ2 tautomer, of the carbonyl oxygen as the reaction progresses from acylenzyme to TI (36). Average distances between the N4 nitrogen and carbonyl oxygen show a general decrease in the TI compared to the acylenzyme, again suggesting that for efficient carbapenemase activity the orientation of the bound carbapenem must be controlled to allow for this increased electrostatic stabilisation of the carbonyl oxygen. Constraining the orientation of bound carbapenem-derived species appears to be the role of the hinge region, which is restricted in its conformational landscape sampled in the simulations of the TI compared to the acylenzyme, resulting in the meropenem-derived species maintaining a conformation that provides greater electrostatic stabilisation to the carbonyl oxygen.

The stability of the deacylating water molecule (DW) is also thought to be an important factor in carbapenemase activity (36, 51, 62). It is logical to infer that enhanced positional stability of a water molecule in the deacylating position positively contributes to efficient deacylation of the meropenem-derived acylenzyme (36, 51). Analysis of the stability of water molecules that sample the DW position suggests that, compared to the other enzymes, BlaC and CTX-M-16 are less well able to stabilise such a water molecule, with the probability of its retention falling below 50 % after less than 200 ps of simulation. This would suggest that activation of the DW, by proton transfer to the Glu166 side chain, is less likely to occur in meropenem acylenzymes of these two enzymes, reducing the frequency of potential nucleophilic attack events required for their deacylation.

To understand the dynamical differences between the acylenzyme and the deacylation tetrahedral intermediate, and their relevance to the rate-determining deacylation step, we analysed the fluctuations (as RMSF values) of individual residues in each simulation. Multiple studies that focus on a single β-lactamase and its variants, and/or reactions with different substrates, have concluded that the stability of the Ω-loop is a factor that is important in determining catalytic efficiency of a given enzyme towards a specific substrate (36, 62–65). Our analyses indicate that, whilst Ω-loop stability is a feature of all four of the carbapenemases investigated here, the acylenzyme ensembles of the carbapenem-inhibited enzymes BlaC and CTX-M-16 both feature conformationally flexible Ω-loops, possibly resulting in catalytically important residues such as Glu166 and Asn170 accessing conformations that are sub-optimal for efficient deacylation. However, visual inspection of the trajectories does not identify any instances during these simulations where Glu166 flips into an ’out’ conformation, as has been observed in simulations of complexes of some class A β-lactamases with other substrates (47, 66). Previous studies have investigated the impact of mutations within the Ω-loop in the KPC-2, TEM-1, PenL and PC1 β-lactamases (36, 62–65). The Ω-loop residue at Ambler position 170 (normally Asn) has also been identified as important for deacylation of carbapenem-derived acylenzymes (19, 62, 67). Increased stability of the Ω-loop will likely promote efficient catalysis, as this increases the likelihood that the Glu166 side chain is positioned optimally for activation of the deacylating water molecule for nucleophilic attack on the carbonyl carbon (C7) of the carbapenem-derived acylenzyme (Figure 3). In simulations of the four carbapenemases considered here, the Ω-loop is stable in both the acyl-enzyme and the TI, indicating that these enzymes are well preorganised for reaction.

For the carbapenem-inhibited enzymes TEM-1 and BlaC, analysis of RMSF values indicates that this loop region undergoes greater fluctuation than is the case for the other enzymes, suggesting that the dynamic ensembles of these enzymes may more regularly sample catalytically incompetent conformations. The carbapenemases, however, have generally more rigid structures, with less flexibility observed in other loop regions, including α3 - α4. This implies that the conformational ensembles of the carbapenemases may be more restricted, sampling a relatively restricted low-energy well upon the potential surface that correlates with a catalytically competent conformation, whilst the carbapenem-inhibited enzymes can more easily sample a shallower, broader area of the free energy surface that includes conformations that do not promote efficient catalysis.

We have previously used dynamical-nonequilibrium molecular dynamics (D-NEMD) simulations, an emerging methodology that identifies intramolecular networks involved in catalysis, to identify the 2β4 (residues 87-94) loop as being important to KPC-2 catalytic activity, validating our findings by kinetic assessment of a KPC-2^G89D^ variant. The simulations here showed that in several of the tested enzymes, including BlaC, CTX-M-16, TEM-1 and SME-1, the 2β4 loop showed an increase in flexibility when progressing from the acyl-enzyme to the tetrahedral intermediate, whilst in SFC-1 this loop decreased in flexibility. This suggests the 2β4 loop may be important in modulating catalysis across class A β-lactamases more generally, and that variants in this loop or screening experiments aiming to identify compounds able to bind this region, may warrant further investigation. Several studies have focused on the importance of transition state stability in promoting efficient catalysis, with catalytically competent enzymes/complexes consistently sampling a low energy well on the transition state, or intermediate, potential energy surface (68, 69). By analysing the per-residue RMSF of both the acylenzyme and TI ensembles, we can compare the stability of different regions in the two states. The analysis appears to indicate that, in all studied enzymes, there is a general increase in stability on transitioning from the acylenzyme to the TI (itself not a true transition state but an intermediate (19, 28)). However, in the carbapenem-inhibited enzymes BlaC, SHV-1 and TEM-1, there appear to be multiple regions of the protein that undergo sizeable decreases in flexibility when comparing the TI to the acylenzyme. This may suggest that, in these enzymes, the ground state (in this case the acylenzyme complex) is not well pre-organised for reaction and is instead able to sample a large area of conformational space, a feature thought to prevent efficient catalysis (70–72).

PCA of the acylenzyme complex and the TI effectively separated enzymes through calculation of overlap or Bhattacharyya coefficients, with carbapenem inhibited enzymes having values of less than 0.25 or 0.5, respectively, and carbapenemases having values above these. Not only does this provide statistical evidence that the carbapenemase enzymes regularly sample transition state-like conformations in their acyl-enzyme ensembles, whilst carbapenem-inhibited enzymes do not, it suggests that a simple computational assay using MD simulations could be used to establish the carbapenemase activity of different class A β-lactamase sequences. With increased compute efficiency and the continued widespread use of protein structure prediction methods, this MD simulation assay potentially provides a method for screening emerging class A β-lactamases for carbapenemase activity.

In conclusion, this investigation has provided broad insights into the dynamic features that promote carbapenemase activity by studying a representative group of class A β lactamases. Our results show that the stabilisation of the acylenzyme carbonyl, through hydrogen bonding to the backbone amides of Ser70 and residue 237, and intramolecular interactions with the acylenzyme amide nitrogen that in turn depend upon orientation of the C3 carboxylate, is a key discriminating factor between the two groups of carbapenem-hydrolysing and carbapenem-inhibited enzymes. This is not however the only criterion that must be fulfilled for efficient carbapenemase activity: carbapenem-hydrolysing enzymes must satisfy multiple requirements for activity, whereas different carbapenem-inhibited enzymes fail in different ways to meet this e.g. by failing to sufficiently stabilise a water molecule in the DW position. Failure in any one requirement is sufficient to prevent efficient catalysis of the deacylation reaction. Furthermore, when comparing the dynamical differences between the acylenzyme and the TI, we show that carbapenemase acylenzymes are dynamically pre-organised for reaction, consistently accessing catalytically competent states, whilst carbapenem-inhibited enzymes do not. Thus, as observed for other enzymes (54, 73–75), changes in the dynamic ensemble of enzyme-bound species as the reaction progresses from a ground state to a transition state or intermediate may be important for catalysis by class A β-lactamases. Understanding the nature of such ensembles, and their relationship to activity, may aid in the development of compounds that could perturb their distribution, either in the acylenzyme or the TI, and so modulate activity of class A β-lactamases towards specific substrates. Furthermore, we demonstrate that similarity between the acylenzyme and TI ensembles discriminates between class A β-lactamases that can and cannot efficiently hydrolyse carbapenems. As bacterial genome sequences become more widely available in clinical microbiology workflows this may find application as part of computational pipelines predicting antibacterial susceptibility from sequence (76).

## Author Contributions

MB conceived the experiments with AJM and JS. MB ran all simulations and analysis. MB wrote the manuscript with inputs from AJM and JS.

## Data Availability

All raw MD simulation data will be made freely available via the University of Bristol’s Research Data Repository (https://data.bris.ac.uk/). Water survival probabilty scripts can be accessed via Github at https://github.com/Michael-Beer/Water-Survival-Probability.git

## Acknowledgements

MB was supported by the BBSRC-funded South West Biosciences Doctoral Training Partnership [BB/T008741/1]. This work is supported by funding to AJM and JS from the European Research Council under the European Horizon 2020 innovation program (PREDACTED Advanced Grant Agreement no. 101021207).

