## Supplementary Information for "A transition state-like acylenzyme conformation distinguishes carbapenemase activity in class A β-lactamases"

^*Corresponding Authors^

***Figure S1 Ca RMSD Analysis of Production MD Simulations of Meropenem Acyl-Enzymes of Class A b-lactamases.*** *The three repeats are plotted separately for the acyl-enzyme MD simulations, with the first frame of each production run used as the reference structure. A) BlaC B) CTX-M-16 C) SHV-1 D) TEM-1 E) KPC-2 F) NMC-A G) SFC-1 H) SME-1. In particular KPC-2 showed higher RMSD changes through the course of the simulations, but these changes related primarily to the increased flexibility of the N- and C-termini compared to the other enzymes.*

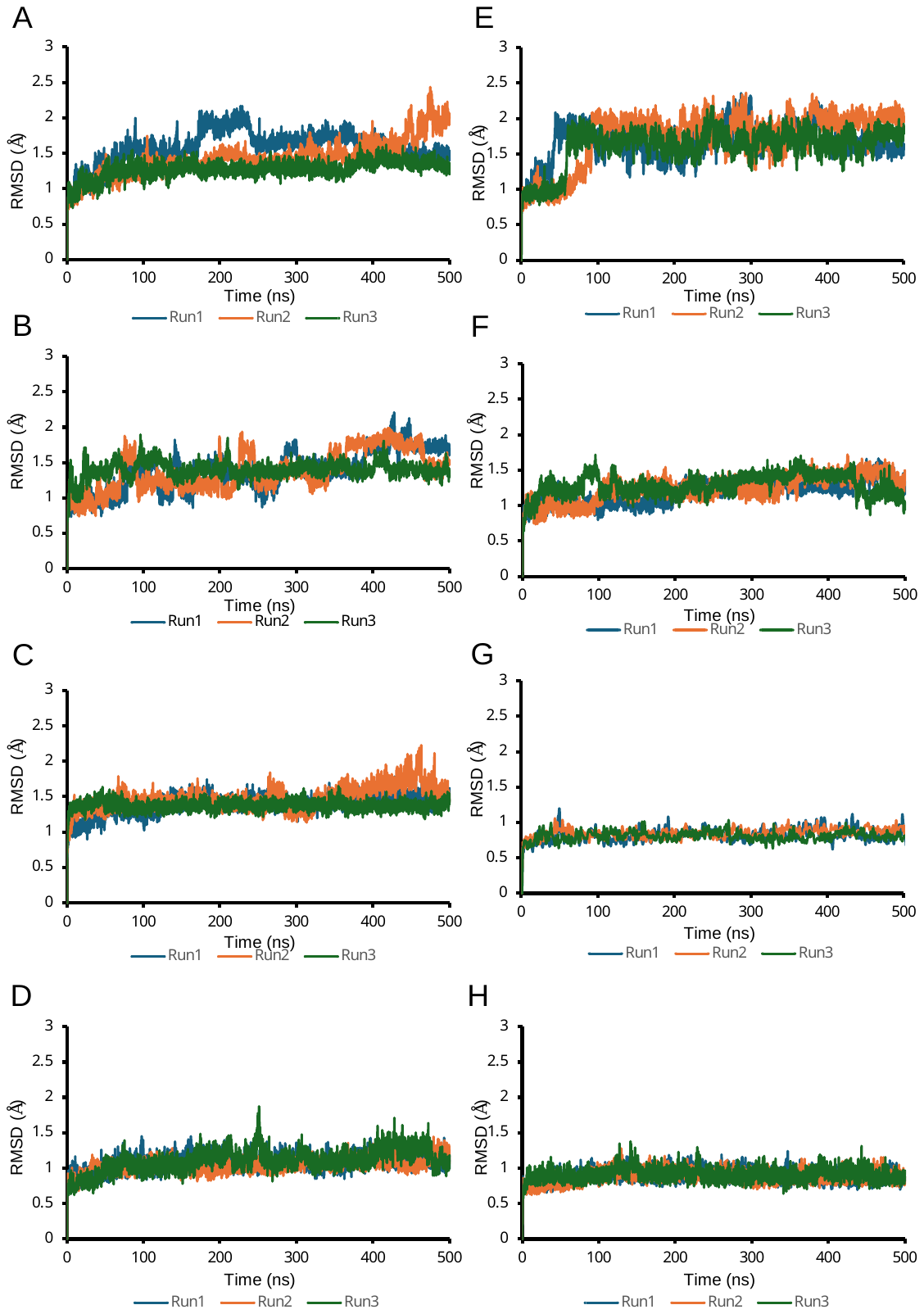

**BlaC**

**CTX-M-16**

**SHV-1**

**TEM-1**

**KPC-2**

**NMC-A**

**SFC-1**

**SME-1**

***Figure S2 Ca RMSD Analysis of Production MD Simulations of Meropenem Tetrahedral Deacylation Intermediates of Class A b-lactamases.*** *A) BlaC B) CTX-M-16 C) SHV-1 D) TEM-1 E) KPC-2 F) NMC-A G) SFC-1 H) SME-1. The first frame of each run was used as the reference structure. The N- and C-termini of the KPC-2 complex appear more flexible than for the other systems and so explain the increased RMSD in the KPC-2 simulations (E).*

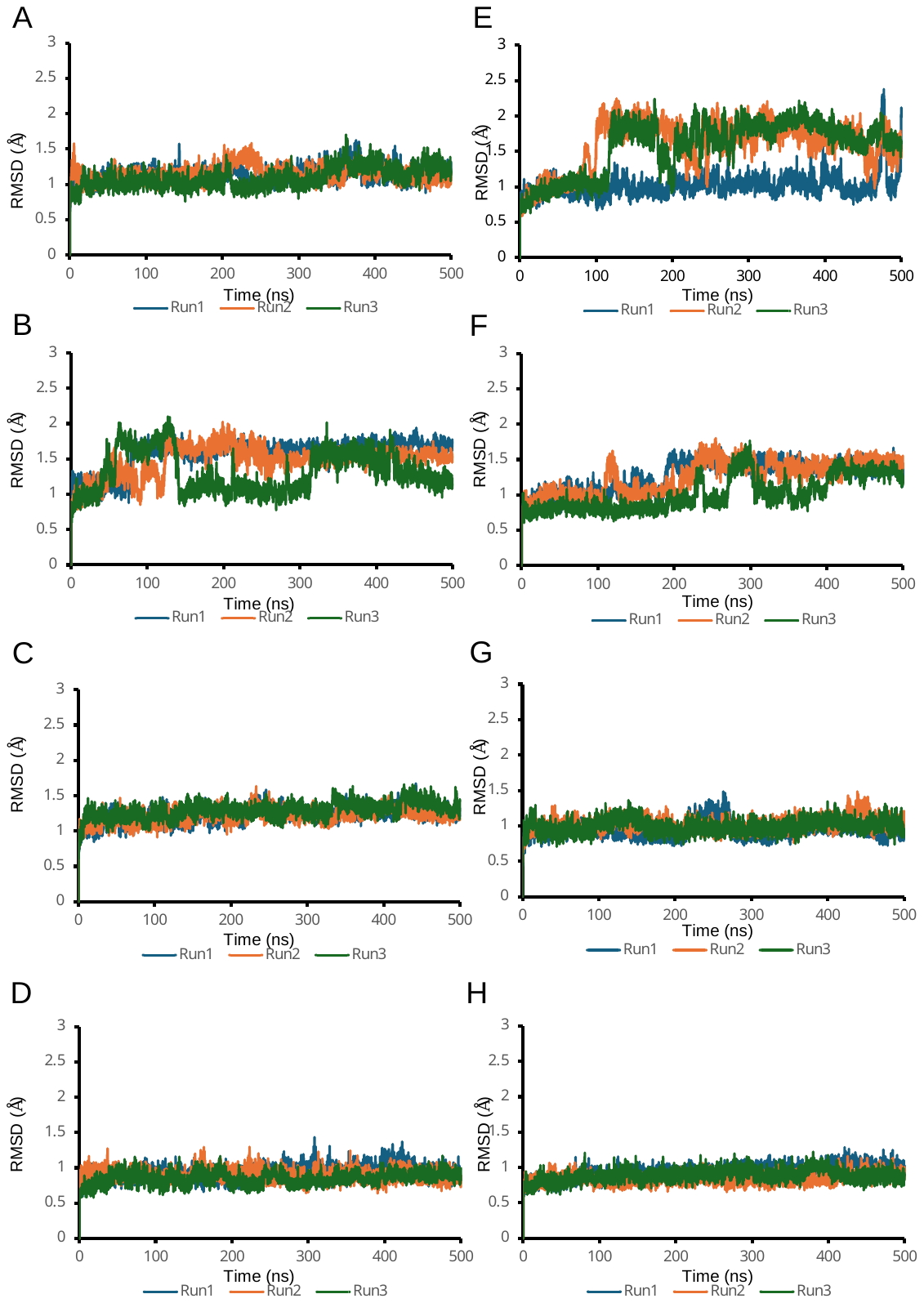

**BlaC**

**CTX-M-16**

**SHV-1**

**TEM-1**

**KPC-2**

**NMC-A**

**SFC-1**

**SME-1**

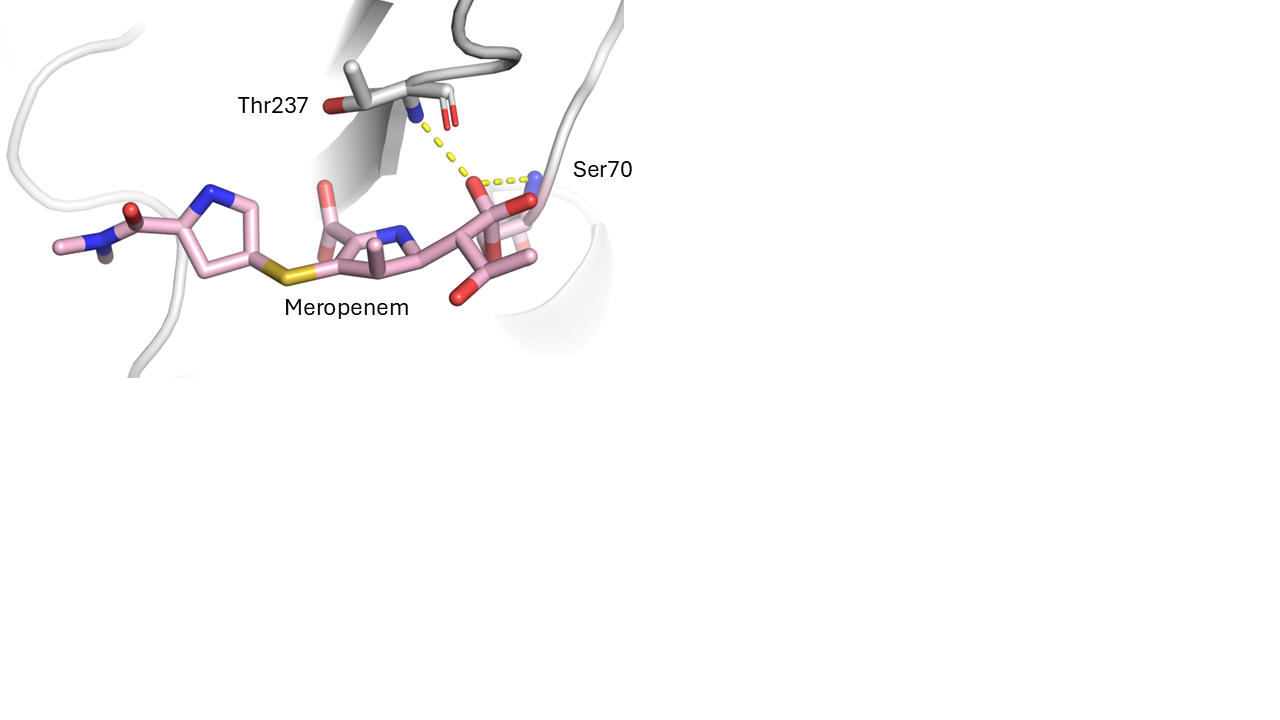

***Figure S3 Meropenem Oxyanion Hole in the Tetrahedral Intermediate (TI).*** *The oxyanion is hydrogen bonded by the backbone amides of residue 237 (threonine in KPC-2, shown here) and Ser70.*

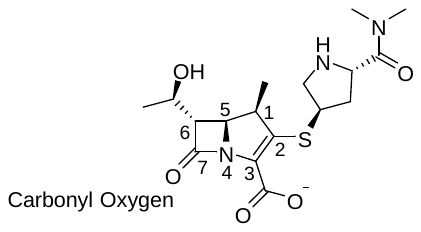

***Figure S4 Meropenem Atom Numbering.*** *The carbonyl oxygen, which forms the oxyanion during the deacylation reaction is also labelled.*

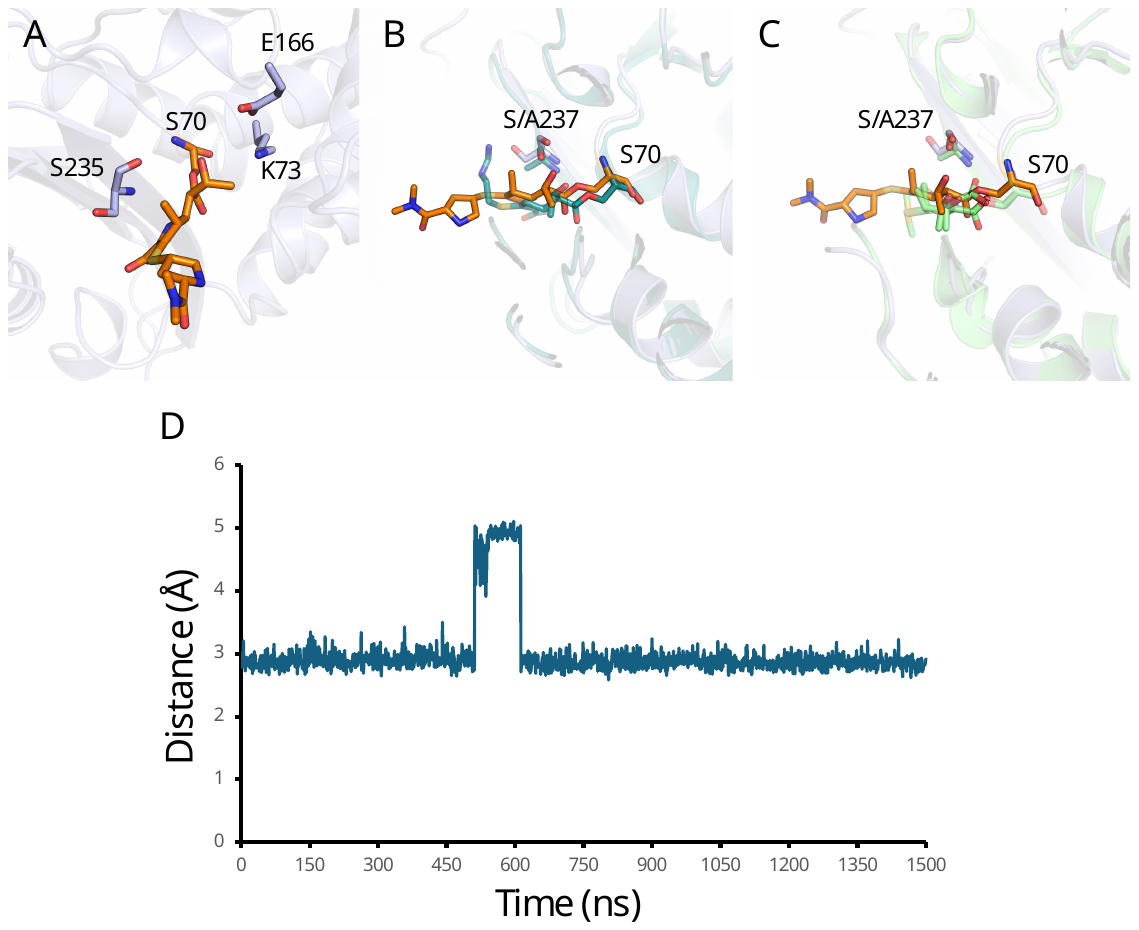

***Figure S5 Carbonyl Oxygen Flipping in CTX-M-16 Acyl-Enzyme Simulations.*** *A) a representative frame where the carboxyl oxygen has flipped out of the oxyanion hole and is not within hydrogen bonding distance of either the backbone amides of Ser70 or Ser237. This is compared to the B) TEM-1:imipenem (PDB ID 2ZD8) and C) SHV-1:meropenem (PDB ID 1BT5) structures in which a flipped orientation of the carbonyl is observed crystallographically. In the SHV-1:meropenem structure two carbonyl oxygen orientations are modelled, one within the oxyanion hole and the other not. D) Distance analysis between the carbonyl oxygen and the backbone amide of Ser70 across the 1.5 μs of simulation (3x500 ns simulations) of the meropenem derived CTX-M-16 acyl-enzyme.*

***Figure S6 Sequence Alignment of the Eight Tested β-Lactamases.*** *Sequences were aligned using the EMBL-EBI Clustal Omega web server. (Note: residue numbers do not correspond to the Ambler numbering scheme as the Ambler numbering scheme excludes the signal sequence (the length of which is variable between different proteins). Figure made in ESPrint 3.0 (1).*

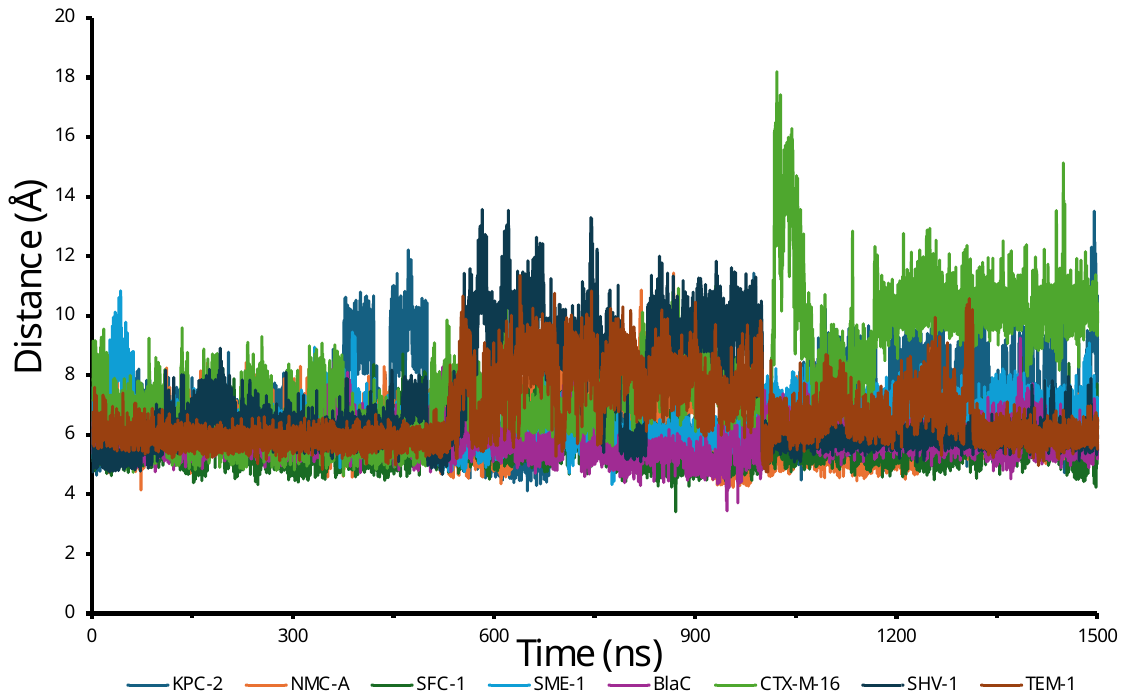

***Figure S7 The Distance between the Hinge Region and the Meropenem Derived Acyl-Adduct.*** *This distance was calculated by measuring between the Cα of residue 216 and the C3 carbon of the meropenem derived acyl-adduct across 1.5 μs of acyl-enzyme simulation.*

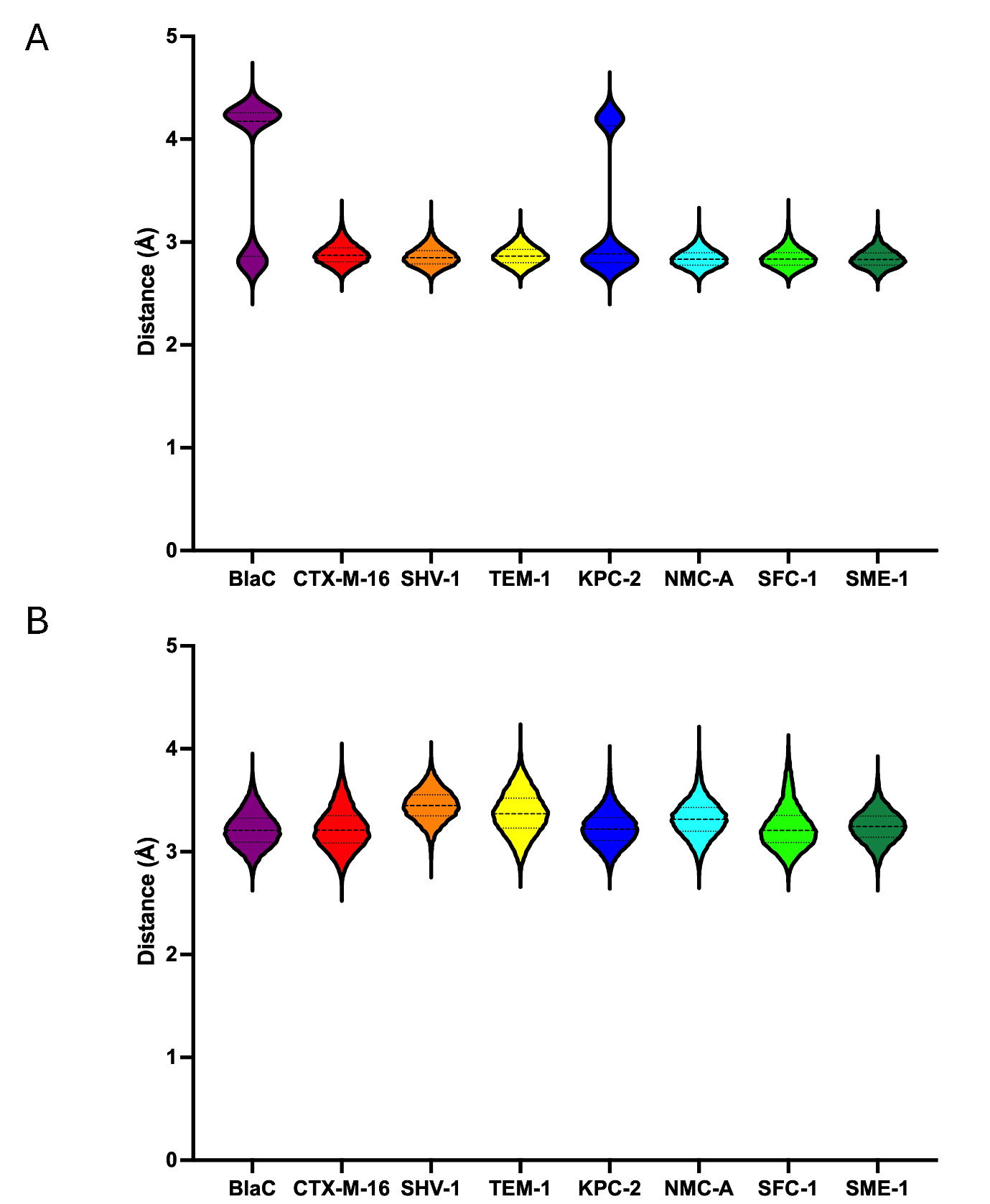

***Figure S8 Average Distance between the β-lactam Amide Nitrogen and Carbonyl Oxygen.*** *The average distance in the acyl-enzyme (A) and the TI (B) is show for each enzyme. ANOVA analysis with pairwise student t-tests indicate that SHV-1 and TEM-1 have a statistically significant greater distance than any carbapenemase in the acyl-enzyme simulations, and BlaC has a statistically significantly greater distance than any carbapenemase in the TI simulations.*

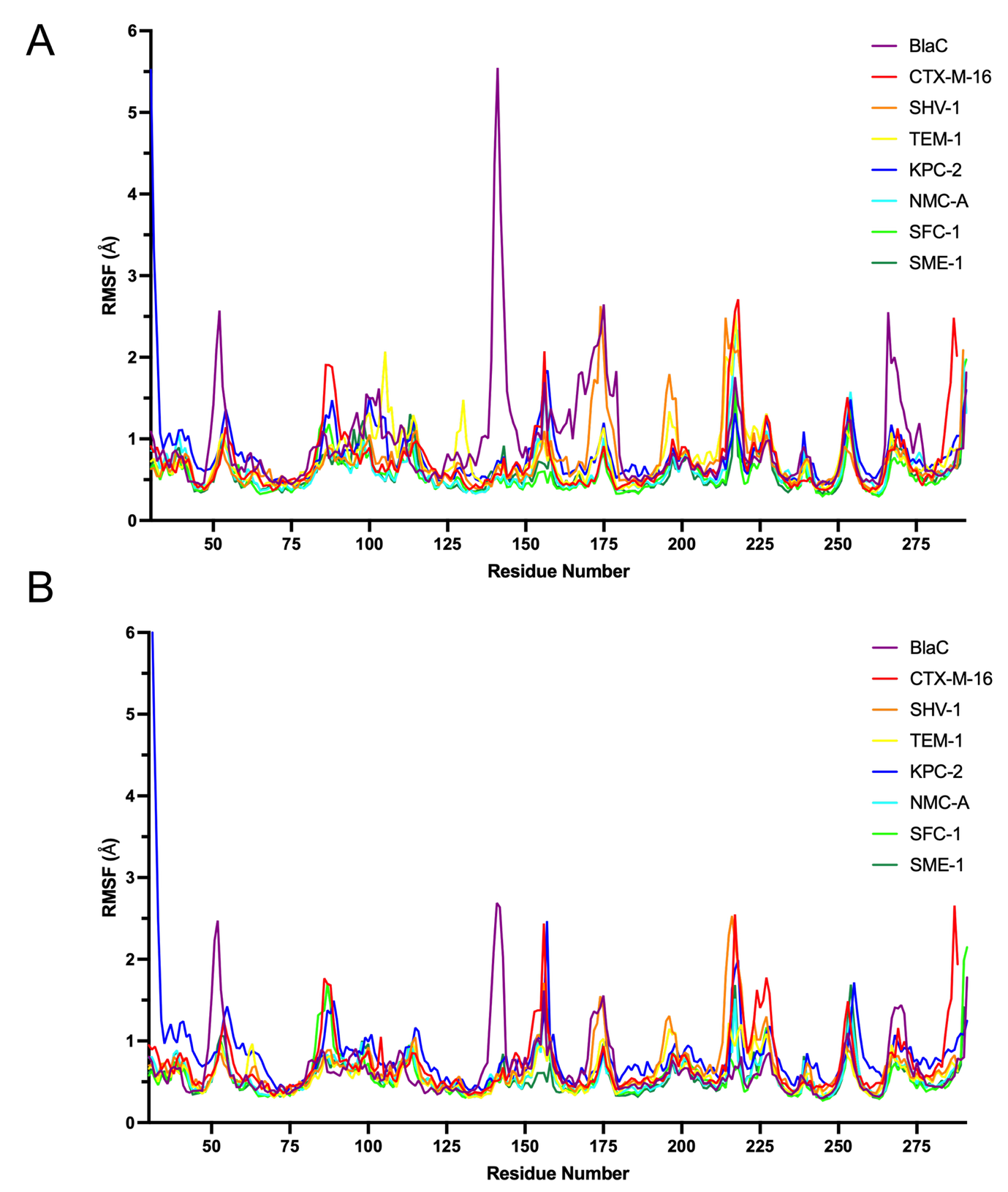

***Figure S9: Per-residue RMSF.*** *Graphs shown for the A) acylenzyme complexes and B) tetrahedral intermediate*

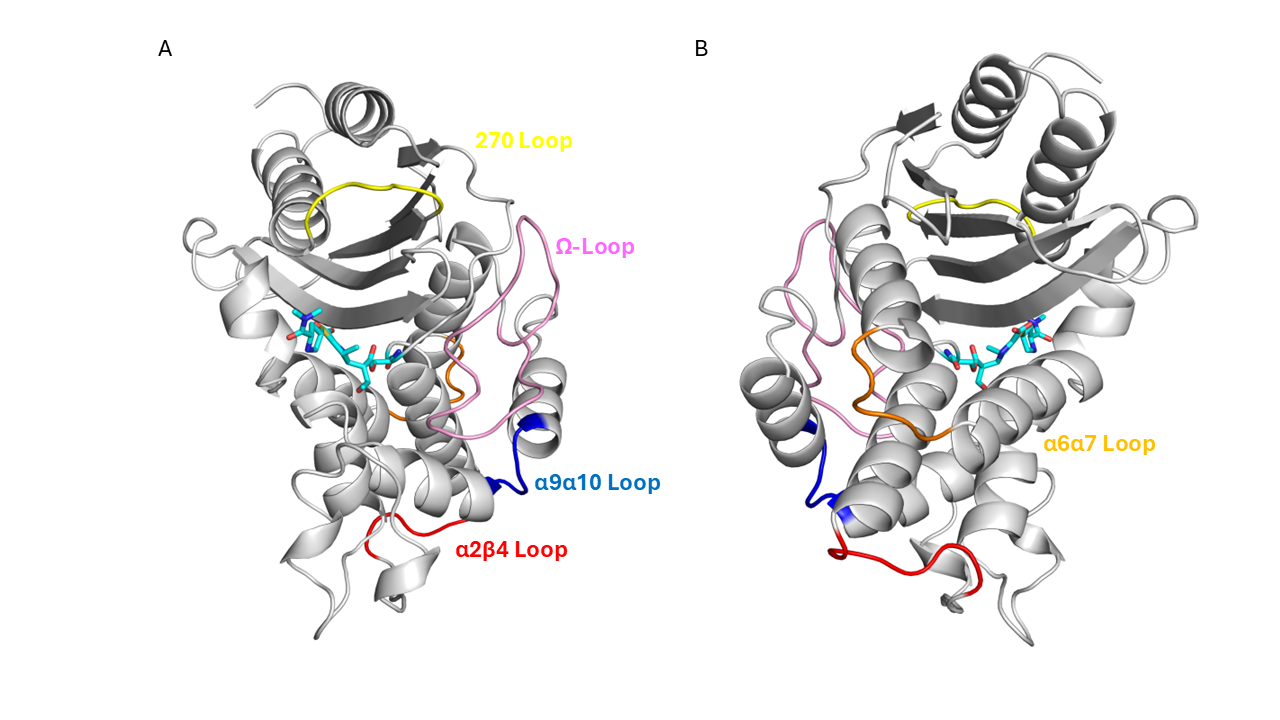

***Figure S10 Loop Regions with High RMSF Scores in the tested β-lactamases.*** *A representative β-lactamase (SHV-1) is shown in A) forward and B) reverse orientations. Flexible loops that have high RMSF values, are shown: α2β4 loop (red), α6α7 loop (orange), Ω-loop (pink), α9α10 loop (blue), 220 loop (purple) and 270 loop (yellow) are shown. Meropenem is also displayed (tan).*

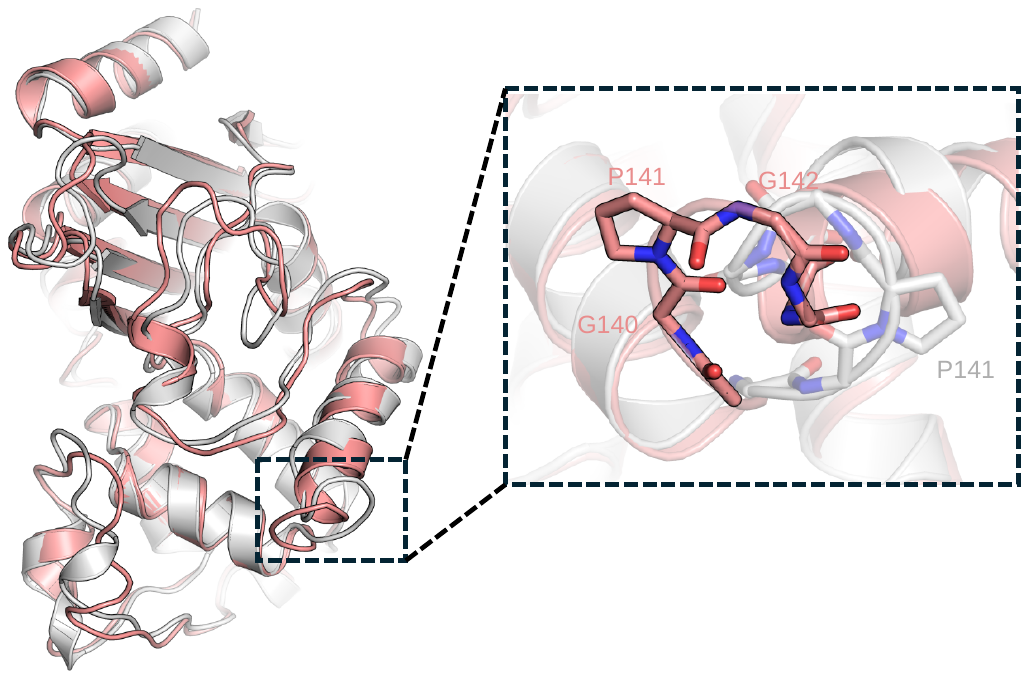
***Figure S11 The Region of High RMSF in BlaC simulations.*** *Structural alignment of a representative from of BlaC from the production MM MD simulations (pink) aligned to the starting structure of BlaC (grey). The α7α6 helix (residues 139-144) was greater in BlaC than any other regions in any of the enzymes (excluding the N- and C-termini). This is large value was contributed to by the movement of G140, P141 and G142. This loops fluctuates between the orientations observed in the representative frame and starting structure conformations.*

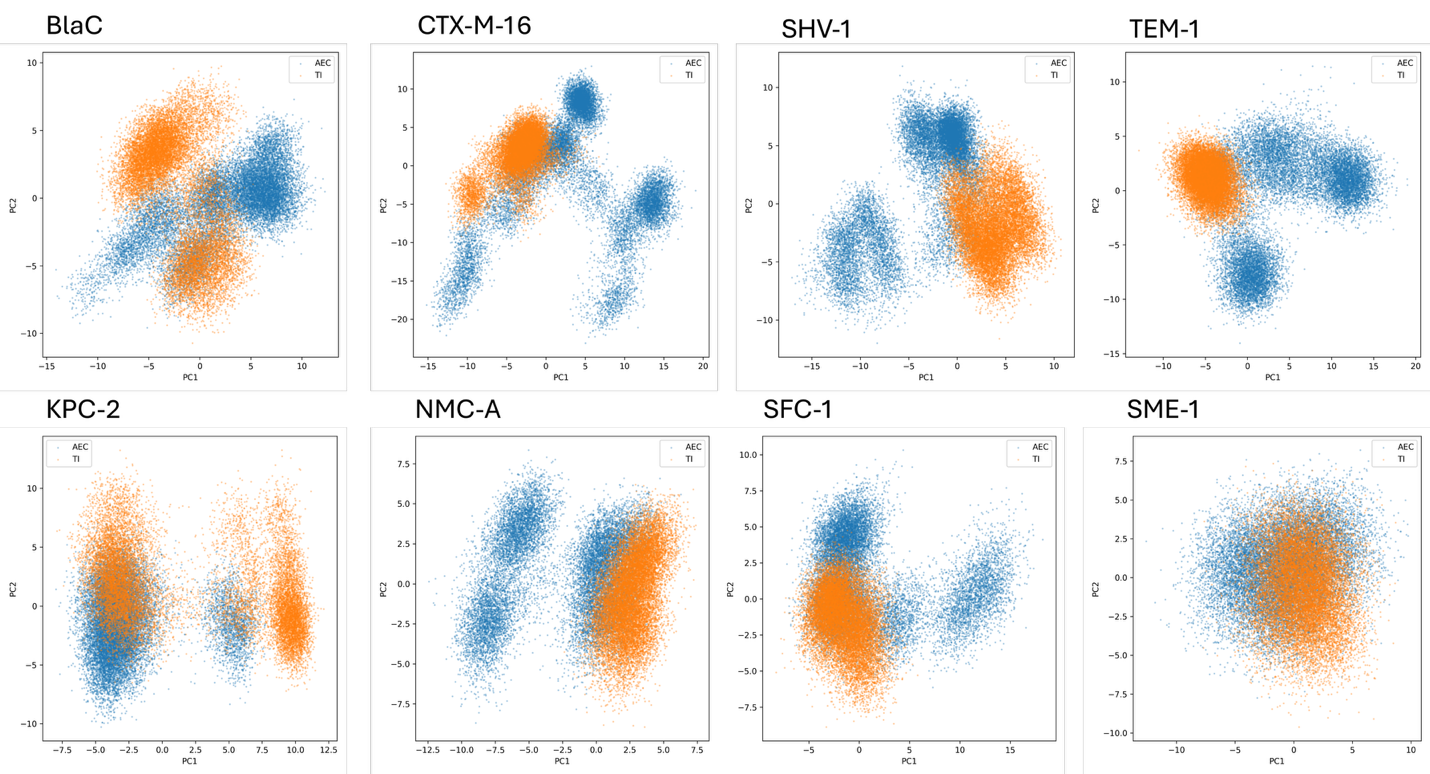
***Figure S12: Principal component analysis (PCA) plots of C⍺ position.*** *The acyl-enzyme complex is shown in blue and the TI in orange, with both projected on the same plane and referenced against the first frame of the first repeat of the AEC simulations.*

***Table S1: Hydrogen bonding percentages throughout the respective acyl-enzyme trajectories.*** *The percentage of simulation frames is shown for different hydrogen bonding interactions between the C3 carboxylate oxygens (Figure S3) and active site residues in the acyl-enzyme. Clear boxes indicates that the hydrogen bonding percentage was less than 10%.*

|  | Ser130 | | Arg220 | | Residue 235 | | Residue 237 | | Lys234 | | Arg244 | | | Arg276 | |
| --- | --- | --- | --- | --- | --- | --- | --- | --- | --- | --- | --- | --- | --- | --- | --- |
| Enzyme | O1 | O2 | O1 | O2 | O1 | O2 | O1 | O2 | O1 | O2 | O1 | O2 | O1 | | O2 |
| BlaC |  |  | 69.7 | 66.2 | 32.1 |  | 13.7 |  |  |  |  |  |  | |  |
| CTX-M-16 |  | 30.4 |  |  |  |  |  |  |  |  |  |  | 64.5 | | 65.5 |
| SHV-1 |  |  |  |  | 69.4 | 40.4 |  |  |  | 17.8 | 52.7 |  |  | |  |
| TEM-1 |  |  |  |  | 56.2 | 33.1 |  |  | 12.7 |  | 22.4 | 49.6 |  | |  |
| KPC-2 |  |  |  |  | 16 |  | 58.3 |  |  |  |  |  |  | |  |
| NMC-A |  |  | 15.5 | 11.4 | 58 |  |  |  |  |  |  |  |  | |  |
| SFC-1 |  |  |  |  | 66.7 |  | 25.5 | 26.7 |  |  |  |  |  | |  |
| SME-1 |  |  | 94.4 | 73.8 |  |  |  |  |  |  |  |  |  | |  |

***Table S2: Hydrogen bonding percentages throughout the respective TI trajectories.*** *The percentage of simulation frames is shown for the hydrogen bonding interactions, calculated by CPPTRAJ, between the C3 carboxylate oxygens (Figure S3) and active site residues in the TI. Red with no numbering indicates that the hydrogen bonding percentage was less than 1%.*

|  | Ser130 | | Arg220 | | Residue 235 | | Residue 237 | | Lys234 | | Arg244 | | | Arg276 | |
| --- | --- | --- | --- | --- | --- | --- | --- | --- | --- | --- | --- | --- | --- | --- | --- |
| Enzyme | O1 | O2 | O1 | O2 | O1 | O2 | O1 | O2 | O1 | O2 | O1 | O2 | O1 | | O2 |
| BlaC |  |  | 100 | 61.8 |  | 90.9 |  | 35.1 |  |  |  |  |  | |  |
| CTX-M-16 |  |  |  |  | 84.3 |  |  |  |  |  |  |  | 94.9 | | 58 |
| SHV-1 |  |  |  |  | 98.6 |  |  |  |  |  | 65.8 | 51.1 |  | |  |
| TEM-1 |  |  |  |  | 98.8 |  |  |  |  |  | 77.7 | 31.8 |  | |  |
| KPC-2 |  |  |  |  |  | 87.3 |  | 89.5 |  |  |  |  |  | |  |
| NMC-A |  |  | 44.2 | 40.7 |  | 65.9 |  | 52.7 |  |  |  |  |  | |  |
| SFC-1 |  |  |  |  |  | 97.4 | 40.2 | 41.2 |  |  |  |  |  | |  |
| SME-1 |  |  | 66.7 | 63.1 |  | 72.4 |  | 24.3 |  |  |  |  |  | |  |

***Table S3: Average distance between the C*⍺ *atoms of residue 69 and residue 238.*** *These two residues form the active site disulfide in* *carbapenemase enzymes. This disulphide is absent in carbapenem-inhibited enzymes due to substitutions in one of these positions. Standard deviation shown in parentheses.*

| ***Enzyme*** | ***Residue 69 – Residue 238 Distance (Å)*** |
| --- | --- |
| *BlaC* | 6.5 (0.4) |
| *CTX-M-16* | 6.1 (0.3) |
| *SHV-1* | 6.3 (0.3) |
| *TEM-1* | 6.2 (0.3) |
| *KPC-2* | 5.4 (0.2) |
| *NMC-A* | 5.2 (0.3) |
| *SFC-1* | 5.4 (0.2) |
| *SME-1* | 5.2 (0.2) |

***Table S4: Overlap and Bhattacharyya coefficients comparing the AEC and TI PCA projections for each enzyme.***

| **Enzyme** | **Overlap Coefficient** | **Bhattacharyya Coefficient** |
| --- | --- | --- |
| BlaC | 0.202 | 0.431 |
| CTX-M-16 | 0.249 | 0.405 |
| SHV-1 | 0.108 | 0.263 |
| TEM-1 | 0.077 | 0.237 |
| KPC-2 | 0.403 | 0.621 |
| NMC-A | 0.273 | 0.513 |
| SFC-1 | 0.370 | 0.630 |
| SME-1 | 0.631 | 0.862 |

***Table S5: Cluster analysis of the AEC and TI PCA projections for each enzyme.*** *Orange indicates the most populous cluster in the AEC projection and green in the TI projection.*

| **Enzyme** | **Cluster 0 (%)** | | **Cluster 1 (%)** | | **Cluster 2 (%)** | |
| --- | --- | --- | --- | --- | --- | --- |
|  | **AEC** | **TI** | **AEC** | **TI** | **AEC** | **TI** |
| BlaC | 68.9 | 100 | 31.1 | 0.0 | - | - |
| CTX-M-16 | 33.2 | 31.9 | 5.5 | 62.3 | 61.3 | 5.8 |
| SHV-1 | 62.1 | 7.2 | 8.2 | 92.8 | 29.7 | 0 |
| TEM-1 | 50.8 | 0.0 | 32.2 | 0.3 | 17 | 99.6 |
| KPC-2 | 11.9 | 46.7 | 88.1 | 53.3 | - | - |
| NMC-A | 53.2 | 100 | 46.8 | 0.0 | - | - |
| SFC-1 | 78 | 100 | 22 | 0 | - | - |
| SME-1 | 17 | 48.2 | 37.7 | 34.5 | 45.3 | 17.3 |
